# A single dimer of the SARS-CoV-2 N protein can associate with multiple fragments of single-stranded and stem-loop RNAs

**DOI:** 10.1101/2025.01.23.634602

**Authors:** Naoya Kaneda, Leo Suzuki, Shun Endo, Syamil Muharror Ahsanul Husna, Shion Ishikawa, Hiroto Takahashi, Hiroyuki Oikawa, Yuji Itoh, Satoshi Takahashi

## Abstract

The nucleocapsid (N) protein of SARS-CoV-2 associates with the viral genomic RNA (gRNA) and forms the ribonucleoprotein (RNP) granules. However, the detailed molecular structures of RNP and their formation mechanism are largely unknown. We used circular dichroism (CD) spectroscopy, fluorescence correlation spectroscopy (FCS) and single-molecule Förster resonance energy transfer (sm-FRET) spectroscopy to understand the interaction between the N protein and different structural units of RNA. We chose polyadenylate chains with 40, 30 and 20 bases possessing a single-stranded structure and three stem loops with 50, 41 and 29 bases selected from gRNA, and labeled their 5’ and 3’ ends by Alexa488 and Alexa647, respectively, for the FCS and sm-FRET measurements. We found that the N protein started to bind to the single-stranded RNAs at the concentrations between 10 and 100 nM. The binding of the N protein to one of the stem loops occurred at the concentration less than 10 nM without melting the stem loop. For all the samples, the binding of multiple molecules of the RNA fragments to a single dimer of the N protein was observed. These results demonstrate that the N protein acts as a non-specific binder to both single-stranded and stem-loop structures of RNA, and that the N protein might contract a long RNA chain by bridging its multiple segments. We propose that the RNP granules might be folded by the association of the numerous stem loops of gRNA triggered by the assembly of the N protein.

## Introduction

Severe acute respiratory syndrome coronavirus 2 (SARS-CoV-2) is a betacoronavirus that caused the COVID-19 pandemic^1^. Coronavirus is a family of positive-stranded RNA viruses having the genomic RNA (gRNA) of ∼30 kb length, which is packaged inside of the lipid bilayer envelope having ∼100 nm diameter^2,3,4^. The virus is further constituted of four structural proteins, the spike (S) protein that is anchored on the viral membrane and binds to cell receptors, the nucleocapsid (N) protein that binds to gRNA inside of virus, and the envelope (E) and membrane (M) proteins that are integral of the viral membranes^5,6^. In this investigation, we focus on the interaction between the N protein and short RNA fragments to formulate a basis for understanding of the assembly of gRNA and the N protein into the RNA–protein complex called ribonucleoprotein (RNP) complex^7^.

The N protein is known to bind to gRNA inside of the SARS-CoV-2 viruses and to form an assembly of 35-40 granules of RNP each having a diameter of ∼15 nm. The overall features of RNP inside of the viruses are sometimes referred to as "eggs in a nest" as well as "beads on a string"^4,7^. Each granule of RNP is assumed to be composed of ∼12 molecules of the N protein and ∼800 bases of gRNA^8^. The RNP structures are likely formed autonomously in the host cytosol upon the assembly of gRNA and the N protein. For example, the *in-vivo* cryo-electron microscopy (cryo-EM) investigations visualized the formation of the granular structures upon the transportation of gRNA to cytosol from the inside of double membrane vesicles (DMVs), an organelle produced in the infected cells used as the storage of the newly synthesized gRNA^4,9^. In the *in-vitro* condition, the granular structures reminiscent of those of RNP were reported to form upon the mixing of the purified N protein and RNA^10,11,12^. Furthermore, the *in-situ* conformational sequencing experiments performed for gRNA in intact viruses revealed the abundance of stem loops formed by the short-range pairing as well as the long-range duplexes and the higher-order junctions, suggesting that the specific structures of RNP might be stabilized by the association with the N protein^13,14^. These results indicate that the N protein triggers the folding of gRNA to form the granular particles of the RNP complex.

A strikingly different property of the N protein in relation to the association with RNA was also reported^15^. The *in-vitro* mixing of single-stranded RNA fragments such as polyU and the purified N protein caused the accumulation of the N protein and the RNA fragments, forming droplets having diameters larger than several micrometers^16,17,18,19,20^. The N protein alone can form droplets at the higher concentration and its mixing of the entire as well as parts of gRNA also caused the droplet formation^8,21^. The regions of gRNA having larger content of the single-stranded structure formed more spherical droplets, while other regions having the higher content of the secondary structures formed more solid-like non-spherical aggregates^21^. Thus, the binding of RNA and the N protein also occurs non-specifically causing the droplet formation reminiscent of the phase separation of intrinsically disordered proteins^22,23^.

The N protein seems to control the specific as well as non-specific interactions with RNA and other proteins to perform its different roles in different stages of the viral infection^11,12^. In the early stages of the viral infection, the replication-transcription complex (RTC) is formed upon the association of several non-structural proteins and produces various lengths of viral RNAs including subgenomic mRNA^2,24,25^. The N protein in host cytosol associates with RTC in the manner similar to the phase separation and enhances the activity of RTC^26,27,28^. The phosphorylation of the serine-arginine rich region (described below) of the N protein makes the droplets more liquid like and might facilitate the viral genome processing^11,12^. In the later stages of the viral infection, the N protein is used for the formation of the granular structures of the newly synthesized gRNA, and is likely to aid the encapsulation of the gRNA by the membrane of the endoplasmic reticulum Golgi intermediate compartment (ERGIC)^4,9^. The specific interaction between the N protein on the surface of RNP and the M protein expressed in the ERGIC membrane might trigger the encapsulation process^29,30^. The latter function of the N protein might be specific to an intact protein without the phosphorylation^11,12^. The mechanism by which a single molecule of gRNA is selectively packaged inside of the virus, sometimes referred to as the packaging, is still largely unknown, but is sometimes assumed to be regulated by the N protein^31^.

The 46-kDa N protein possesses two folded domains, the N-terminal domain (NTD) (residue 49 to 174) and the C-terminal domain (CTD) (247 to 364) that are connected by an intrinsically disordered linker (linker) (175–246) and flanked by the disordered N terminal tail (NT-tail) (1–48) and C terminal tail (CT-tail) (365–419)^32,33^. The NTD possesses a basic RNA-binding cleft and binds with RNA^34,35^. The CTD forms a tightly associated dimer, possessing a basic groove that can associate with RNA^32,36^. The NT-tail also binds to RNA and facilitates the RNA binding of NTD^37^. The CT-tail possesses a region that associates with nucleic acids and might contribute for the tetramerization of the N protein^38,39^. The linker region further contains the serine-arginine (SR) rich region (176–209) used as the phosphorylation sites and the leucine-rich sequence (218–231) identified as the domain regulating the higher oligomerization^40^. The complex architecture of the N protein composed of the multiple domains with distinct properties bestows the protein to possess highly heterogeneous and dynamic structures in the isolated state, and an ability to form multiple oligomeric forms and to bind to RNA at the multiple sites, causing its complex behaviors upon the association with RNA^19,31,41^.

Many of the important functions of the N protein and gRNA are thus still unresolved due to the complexity of the N protein and gRNA interactions. In this investigation, we aimed to reveal the association state and the structural changes of the short fragments of RNAs having both single-stranded and stem-loop structures upon the binding with the N protein. To investigate the association state of the RNA samples, we utilized fluorescence correlation spectroscopy (FCS). To investigate the structural changes of the RNA samples, we utilized single molecule Förster resonance energy transfer (sm-FRET) spectroscopy based on the alternating laser excitation (ALEX) detection method. We further used the circular dichroism (CD) spectroscopy to investigate the secondary structure content of the RNA samples.

## Results

### Purification of the N protein

We expressed the N protein tagged with His10 at the N terminus in *E. coli* and purified it by using the nickel affinity column. As pointed out previously, a trace amount of contaminating nucleotides could cause an aggregation of the sample^12,42^. To obtain reproducible spectroscopic results, it was important to eliminate the nucleotide contaminations by washing the cell extracts repeatedly in the presence of nuclease. In addition, the nickel column purification should be conducted in the urea unfolded state and the purified sample should be flash frozen in small aliquots in the storage buffer containing 1 M NaCl to avoid repeated freezing and thawing. The details of the sample expression and purification are explained in Supporting Texts. The SDS PAGE results of the purified sample showed that the sample did not contain other protein impurities (Supporting Figure S1A). The UV absorption spectra showed only the absorption without light scattering having the A_260nm_/A_280nm_ ratio of 0.6, showing the absence of nucleotides and aggregated particles (Supporting Figure S1B). The circular dichroism (CD) spectrum of the purified sample showed mostly the negative ellipticities characteristic of the unfolded polypeptides and a small positive feature at around 230 nm consistent with the reported CD spectrum of the N protein^43,42^ (Supporting Texts and Supporting Figure S1C).

### Selection of the short RNA fragments

We chose six fragments of the RNA samples to understand how the basic structural units of RNA behave upon the mixing with the N protein. To understand the changes in the flexible single-stranded RNAs, we chose 40-, 30- and 20-base polyadenylates, rA_40_, rA_30_ and rA_20_. By comparing the results for the three samples, we expected to identify changes dependent on the chain length. We further chose sequences selected from gRNA that were shown to form stem-loop structure. The gRNA contains numerous stem loops whose lengths are typically 60–80 bases. We chose the 4^th^ stem loop from the 5’ end (SL4), corresponding to the bases from 78 to 127 of gRNA. In addition, we chose additional two stem-loop structures, PS-SL1 and PS-SL2, which were hypothesized as the "packaging signal" sequences^44^. The packaging signal is the hypothetical region of gRNA that might interact specifically with the N protein and trigger the packaging of gRNA by associating with the M protein^31^. PS-SL1 and PS-SL2 correspond to the bases from 19915 to 19955 as well as from 19973 to 20001 of gRNA, respectively. The sequences selected were listed in Table 1. For FCS measurements, the 5’ end of the RNA fragment was labeled with Alexa488, and were denoted as 488-rA_40_ etc. For ALEX measurements and in some FCS measurements, the 5’ and 3’ ends of the RNA fragments were labeled with Alexa488 and Alexa647, respectively, and were denoted as 488-rA_40_-647 etc. The samples used for the CD measurements were without the fluorophore labeling.

**Table 1.**
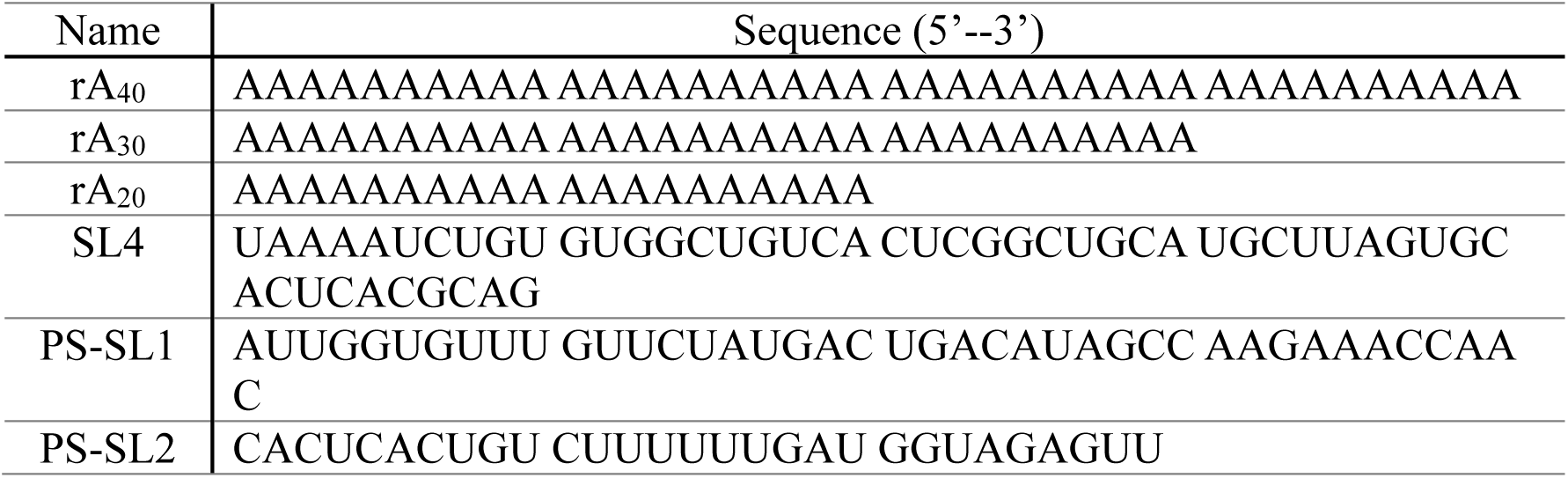
Sequences of the RNA fragments investigated in this study.

### CD spectroscopy of rA_40_ and SL4

To understand the structural changes at the ensemble level, we first observed the CD spectra of the selected RNA samples, rA_40_ and SL4, upon the binding of the N protein. The CD spectrum for rA_40_ shows positive and negative ellipticities at 263 nm and 248 nm, respectively, characteristic of the single-stranded π-stacked helical structures (Figure 1A, red)^45,46,47^. The addition of the N protein reduces the ellipticities slightly, indicating that a small decrease in the helical secondary structure upon the binding of the N protein (Figures 1A, yellow and green). In the case for SL4, the spectrum in the absence of the N protein shows a positive feature at 265 nm (Figure 1B, red), corresponding to the right-handed stem-loop structure^48^. The addition of the N protein to the sample did not change the features at all (Figures 1B, yellow and green), showing that the binding of the N protein did not break the stem loop. We note that the samples did not show any scattering even at the micromolar concentrations of RNAs and the N protein, suggesting the absence of aggregates. Thus, the binding of the N protein might change the structure of the single-stranded RNA slightly but not the stem-loop structure.

**Figure 1.**
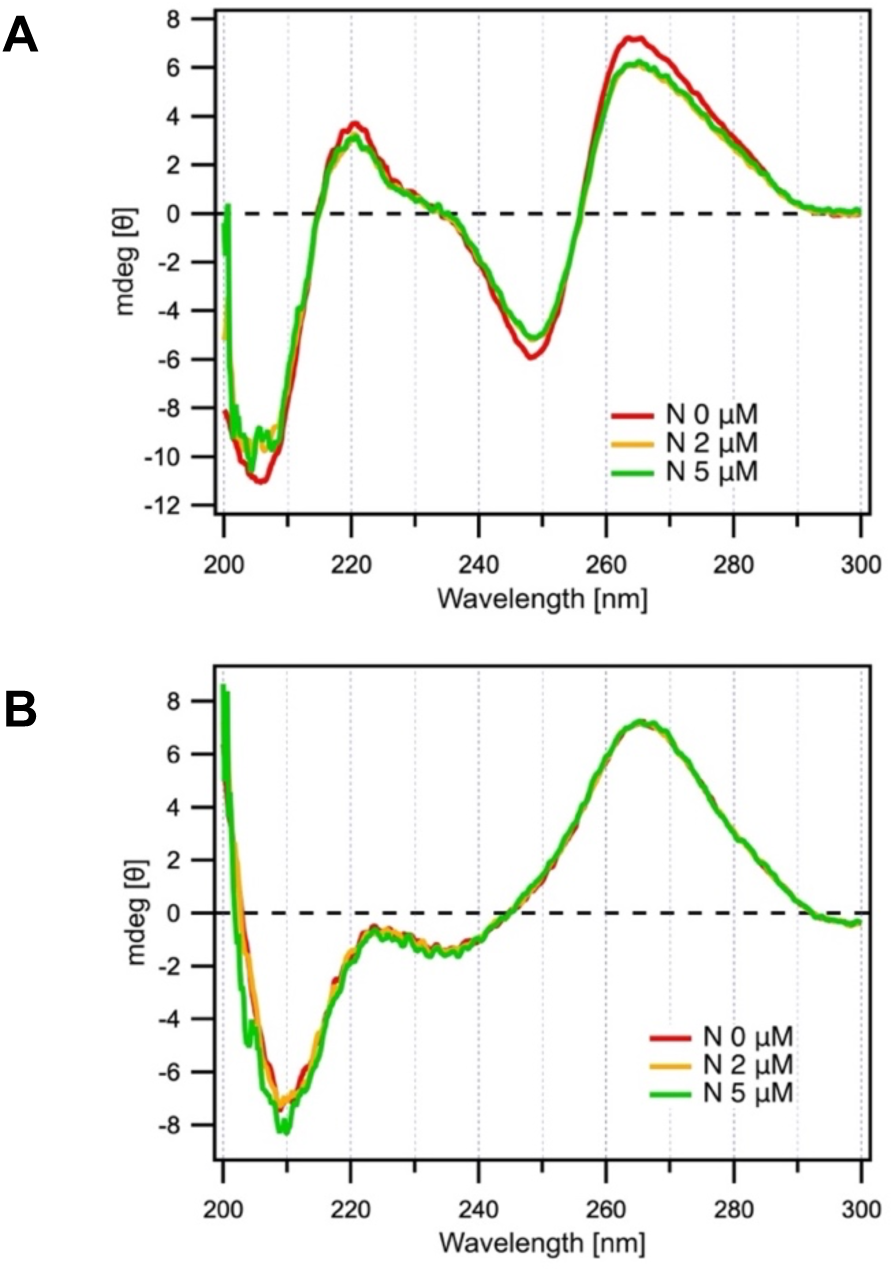
The CD spectra of rA_40_ and SL4 upon the association with the N protein. A) The CD spectra for 2-μM rA_40_ in the absence (red) and presence of the N protein at 2 (yellow) and 5 μM (green). B) The CD spectra for 5-μM SL4 in the absence (red) and presence of the N protein at 2 (yellow) and 5 μM (green). For both panels, the CD contributions of the N protein (Supporting Figure S1C) were subtracted. Details of the CD measurements were explained in Supporting Texts.

### FCS measurements for the singly labeled RNA samples at the 484-nm excitation

To investigate the association state of the RNA samples and the N protein, we next conducted the FCS measurements upon the excitation of Alexa488 at 484 nm. We prepared 1-nM solution of 488-rA_40_, 488-rA_30_, 488-rA_20_ and 488-SL4, and measured their FCS data in the absence and presence of 0.1, 1, 10, 100 and 1000 nM of the N protein. The experimental detail of the FCS measurements were explained in Supporting Texts. The correlation data were presented in Supporting Figures S2. We found that the labeled RNA samples contained non-separatable free fluorophore and that the averaged fluorescence intensity,, became smaller at the higher concentration of the N protein, suggesting the fluorescence quenching of the labeled RNA upon the association of the N protein (Supporting Table S1). Thus, we analyzed the correlations assuming two emitting components having different diffusivity and brightness, one is the free Alexa488 diffusing quickly with a constant brightness and the other is the labeled RNA samples diffusing slowly with a variable brightness. The general expression of the fluorescence correlation for the two-component system with one component having a variable brightness can be described as follows (eq. 1):

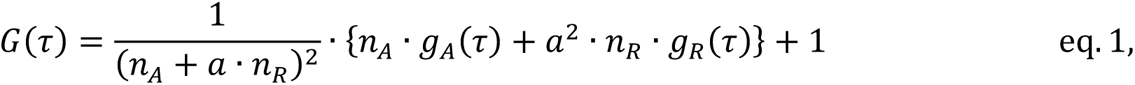

where *n*_A_, *n*_R_ and *a* correspond to the number of free fluorophore in the observation volume, the number of the labeled RNA, and the ratio of the brightness of the labeled RNA relative to that of the free fluorophore, respectively^49,50^. 𝑔_𝐴_(𝜏) and 𝑔_𝑅_(𝜏) are the normalized correlations for the translational diffusion of the free fluorophore having the translational diffusion time 𝜏_𝐴_, and the labeled RNA having the translational diffusion time 𝜏_𝑅_, respectively. To explicitly incorporate the fluorescence intensity information, we conducted the fitting based on the modified equation:

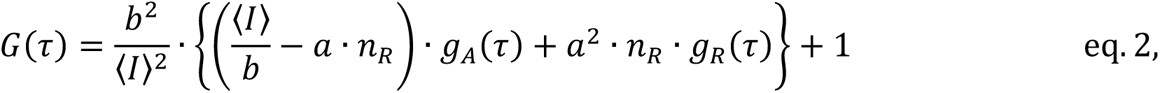

where *b* is the brightness of the free fluorophore. The derivation of eq. 2 from eq. 1 is based on the equality that <*I*> is equivalent to 𝑏 · (𝑛_𝐴_ + 𝑎 · 𝑛_𝑅_). Under the assumption that *a* is 1 for the RNA sample in the absence of the N protein and that *a* might change upon the addition of the N protein, we determined *a*, *n*_R_ and the hydrodynamic radius, *R*_H_, of the labeled RNA sample based on the fitting analysis. Further detail of the analysis was explained in Supporting Texts. The parameters obtained by the fitting were listed in Supporting Table S1. The important parameters were plotted in Figure 2.

**Figure 2.**
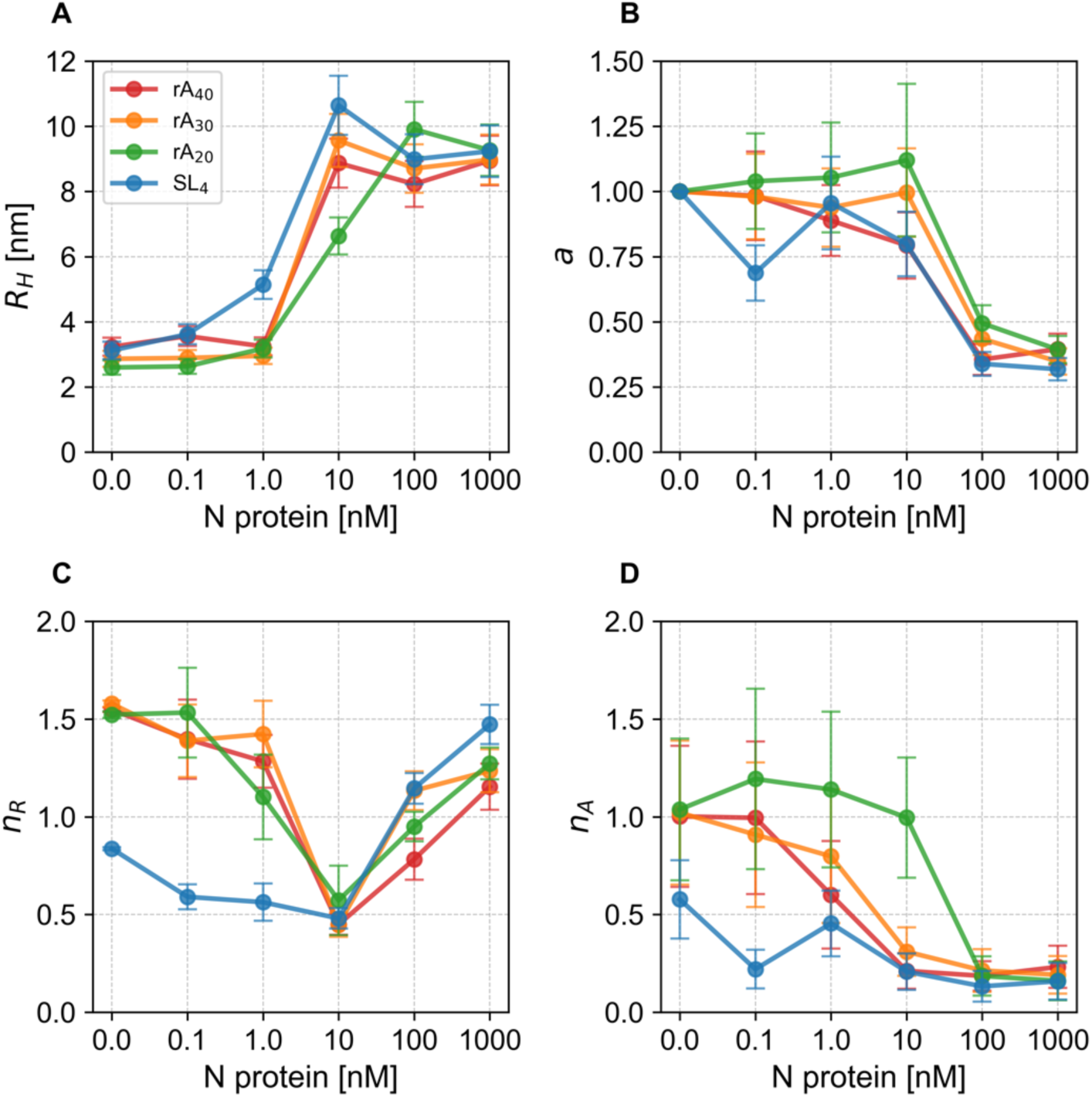
The parameters obtained by the analysis of the FCS data excited at 484 nm for 488-rA40, 488-rA30, 488-rA20 and 488-SL4 at different concentrations of the N protein. The FCS data were analyzed assuming two diffusing components, the free Alexa488 diffusing quickly having constant brightness and the labeled RNA diffusing slowly having variable brightness. For all the panels, the data for 488-rA40, 488-rA30, 488-rA20 and 488-SL4 were shown in red, orange, green and blue, respectively. The error bars were roughly estimated by assuming 10 % changes of the observed fluorescence intensities as explained in the supporting text. A) The hydrodynamic radius, *R*_H_, of the RNA particles. B) The ratio of brightness, *a*, of the RNA particles relative to the brightness of the free Alexa488. C) The number of the RNA particles, *n*_R_, in the observation volume. D) The number of the free Alexa488, *n*_A_, in the observation volume.

We will first examine the results for 488-rA_40_ that suggested the association of multiple RNA molecules to a single dimer of the N protein. *R*_H_ of 488-rA_40_ in the absence and presence of up to 1 nM of the N protein was ∼3.2 nm, corresponding to *R*_H_ of free 488-rA_40_ (Figure 2A, red). The addition of 10 nM of the N protein resulted in the increase of *R*_H_ to ∼8.9 nm, which was constant even at 1000 nM of the N protein. The brightness ratio, *a*, was nearly one at 0.1 nM of the N protein but became slightly smaller (∼0.8) upon the association of the N protein at 10 nM and was further reduced to ∼0.4 at 1000 nM (Figure 2B, red). The results indicate that the association of the N protein quenches the Alexa488 fluorescence of the labeled sample. The number of the labeled RNA molecule in the observation volume, *n*_R_, was 1.3∼1.5 in the absence and presence of the N protein up to the concentration of 1 nM, but drops to ∼0.5 at 10 nM but increase again to ∼1.2 at 1000 nM (Figure 2C, red). Since *R*_H_ indicates that the N protein is bound to 488-rA_40_ at the N protein concentration of 10 nM, the drop of *n*_R_ at 10 nM suggests that multiple RNA molecules are bound to one dimer of the N protein. The increase of *n*_R_ at the concentrations of the N protein higher than 100 nM was likely the result of the association of single RNA to the N protein dimer due the abundance of the latter over the former.

In the case for other RNA samples, the estimated parameters showed behaviors similar to those observed for 488-rA_40_, and supported the association of multiple RNA molecules to one dimer of the N protein at 10 nM. *R*_H_ of 488-rA_30_ showed the sigmoidal change similar to 488-rA_40_, demonstrating the binding of the N protein at around 10 nM (Figure 2A, orange). The association of 488-rA_20_ and the N protein was not completed at 10 nM (Figure 2A, green). In contrast, 488-SL4 started to associate with the N protein at 1 nM (Figure 2A, blue). The reduction of *a* at the N protein concentrations higher than 100 nM was similarly observed for all the samples, again confirming the quenching of the Alexa fluorescence by the association of the N protein (Figure 2B). The *a* value for 488-A_30_ and 488-A_20_ became slightly large at 10 nM, suggesting the binding of multiple RNA fragments having partially quenched fluorescence to one dimer of the N protein. Most notably, the *n*_R_ values for all the samples were smallest at 10 nM but increased at 100 nM and 1000 nM, showing again that the multiple fragments bound to one dimer of the N protein at 10 nM (Figure 2C).

In summary, the FCS measurements for the RNA samples labeled with Alexa488 demonstrated that the binding of the RNA and the N protein occurs at the very low concentration range. Among the single-stranded RNAs, the longer samples showed the higher affinity having the dissociation constant near 10 nM. In contrast, the double-stranded SL4 likely possessed the dissociation constant smaller than 10 nM. The *n*_R_ value became always the smallest at 10 nM of the N protein, suggesting the association of two or more RNA molecules to a single dimer of the N protein. We should note that the number of the free fluorophore, *n*_A_, changed upon the increase of the N protein concentration (Figure 2D). We confirmed separately that the free Alexa488 does not directly associate with the N protein (not shown). We suggest that the results might be caused by a possible contamination of Alexa488-labeled RNA having much smaller number of bases. However, the major conclusion that two or more RNA molecules might associate with a single dimer of the N protein does not change by this possible presence of an impurity.

### FCS measurements for the doubly labeled RNA samples at the 642-nm excitation

To confirm the association of multiple RNA molecules to a dimer of the N protein, we next conducted the FCS measurements for the doubly labeled samples, 488-rA_40_-647 and 488-SL4-647, upon the excitation of Alexa647 at 642 nm, since the quenching efficiency of the fluorophore by the binding of the N protein might not occur for Alexa647. For 1 nM and 0.5 nM of 488-rA_40_-647 and 488-SL4-647, respectively, we added 0, 10 and 100 nM of the N protein, and conducted the FCS measurements excited at 642-nm using the observation system different from that used for the 484-nm excitation (Supporting Texts). We analyzed data assuming the two diffusing components as we analyzed the FCS data at the 484-nm excitation. The correlation data for 488-rA_40_-647 and 488-SL4-647 were presented in Supporting Figures S3A and B, respectively. The fluorescence intensity data as well as the fitted parameters were listed in Supporting Tables S2. The important parameters were plotted in Figure 3.

**Figure 3.**
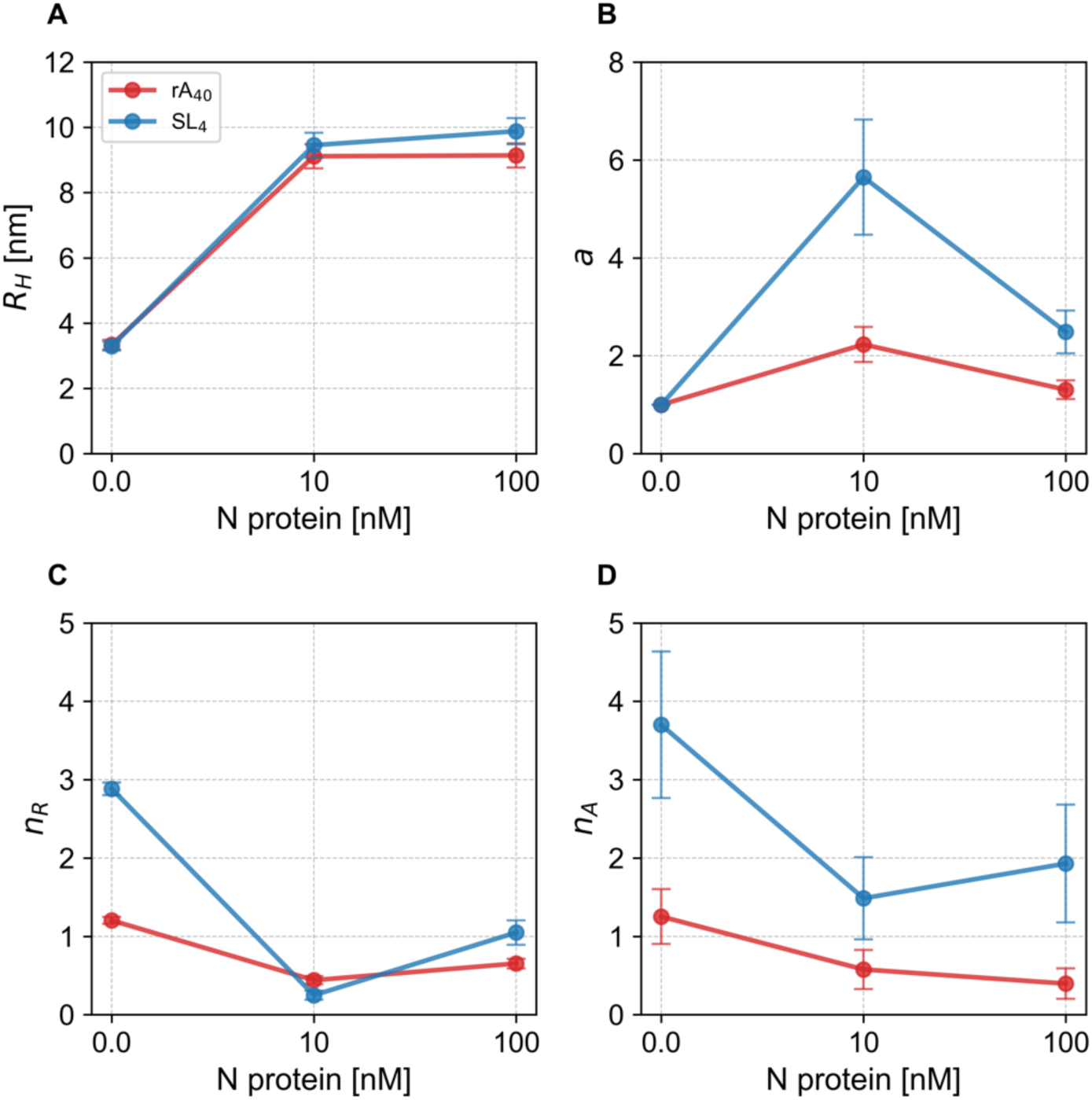
The parameters obtained by the analysis of the FCS data excited at 642 nm for 488-rA_40_-647 and 488-SL4-647 at different concentrations of the N protein. The FCS data were analyzed assuming two diffusing components, the free Alexa647 and the labeled RNA. For all the panels, the data for 488-rA_40_-647 and 488-SL4-647 were shown in red and blue, respectively. A) The hydrodynamic radius, *R*_H_, of the RNA particles. B) The ratio of brightness, *a*, of the RNA particles relative to the brightness of the free Alexa488. C) The number of the RNA particles, *n*_R_, in the observation volume. D) The number of the free Alexa488, *n*_A_, in the observation volume.

In the case for 488-rA_40_-647, the addition of the N protein at 10 nM increased *R*_H_ from the free RNA value of 3.3 nm to 9.1 nm, both of which are consistent with the values estimated by the FCS measurements excited at 484 nm (Figure 3A, red). However, *a* of 488-rA_40_-647 complexed with the N protein at 10 nM became ∼2.2 but returned ∼1.3 at 100 nM (Figure 3B, red). The *n*_R_ value was ∼1.2, ∼0.4 and ∼0.7 at the N protein concentrations of 0, 10 and 100 nM, respectively (Figure 3C, red). These results can be consistently explained by the association of two or more molecules of 488-rA_40_-647 to a single dimer of the N protein at 10 nM, and by the association of a single molecule of 488-rA_40_-647 to a single dimer of the N protein at 100 nM. While the data were rather scattered, the increase in *a* and the decrease in *n*_R_ only at 10 nM of the N protein were similarly confirmed for SL4 (Figure 3B, blue). The *a* value for SL4 at 10 nM was more than 5 and the *n*_R_ value was less than 25 % of that in the absence of the N protein, further suggesting multiple molecules of SL4 might bind to the N protein dimer at 10 nM. Thus, the FCS data at the 642 nm excitation supported that the association of multiple RNA fragments to one dimer of the N protein at 10 nM.

The raw data of the fluorescence intensity fluctuation obtained at the 642-nm excitation of 488-rA_40_-647 and 488-SL4-647 showed frequent "spikes" only in the presence of the N protein, which suggested an appearance of particles having larger numbers of the fluorophores (Supporting Figure S4). The non-stational fluctuation of the fluorescence signals such as spikes causes oscillating features in the correlograms as demonstrated in the negative fitting residuals of the correlograms in the longer correlation time (Supporting Figure S3). In the correlation data obtained at the 484-nm excitation, we did not observe such spikes likely because of the quenching of the Alexa488 fluorescence. Thus, our data demonstrated that a small fraction of the samples forms larger aggregates.

### sm-FRET measurements of the doubly-labelled rA_40_, rA_30_ and rA_20_

To investigate structural changes of the RNA samples upon the association with the N protein, we next conducted the sm-FRET measurements of the doubly-labeled RNA samples at different concentrations of the N protein. We utilized the ALEX measurement system, in which two lasers excite donor and acceptor fluorophores alternatively to obtain the FRET efficiency (*E*) and the donor and acceptor stoichiometry (*S*), suggesting the changes in the distance between the donor and acceptor and in the molar ratio of the donor and acceptor contained in one burst, respectively. We further analyzed the burst width and brightness which correspond to the residence time of a fluorescent particle in the focus volume, and the rate of the photon emission of the particle determined by the number of photons divided by the burst width, respectively. As explained in Supporting Texts, we established the sample preparation protocol and the burst selection condition that could reproduce the experimental results.

We first examined the addition of the N protein to the dilute solutions of 488-rA_40_-647, 488-rA_30_-647 and 488-rA_20_-647, and showed their FRET efficiency probability distributions in Figure 4A. The peak FRET efficiency is less than 0.1 for 488-rA_40_-647 and 488-rA_30_-647, and ∼0.2 for 488-rA_20_-647. The values are consistent with the extended helical configuration of polyadenylate samples^45–47^. The addition of the N protein increased the peak FRET efficiency to ∼0.3, ∼0.4 and more than 0.5 for 488-rA_40_-647, 488-rA_30_-647 and 488-rA_20_-647, respectively, and broadened the distributions ranging from 0 to 1. While this result might suggest a heterogeneous reduction of the donor-acceptor distance, the data might also be interpreted by the quenching of the donor fluorophore and by a plausible fixation of the rotational freedom of the labeled fluorophores. The addition of the N protein caused the changes in the stoichiometry distribution of the RNA samples (Figure 4B). For all the three samples, the stoichiometry in the absence of the N protein was peaked at ∼0.3, which corresponds to the samples possessing Alexa488 and Alexa647 at the 1:1 ratio (Supporting Texts). The stoichiometry became smaller and partially shifted to ∼0.2 in the presence of the N protein. This is consistent with the quenching of Alexa488 fluorescence by the binding of the N protein. The binding of the N protein also caused the expansion of the burst width (Figure 4C). In the absence of the N protein, the burst width distribution was peaked at ∼5 ms and possessed a narrow width, reflecting the fast diffusion. However, the addition of the N protein shifted the peak to ∼7 ms and broadened the distribution, clearly demonstrating the binding of the N protein to the RNA samples.

**Figure 4.**
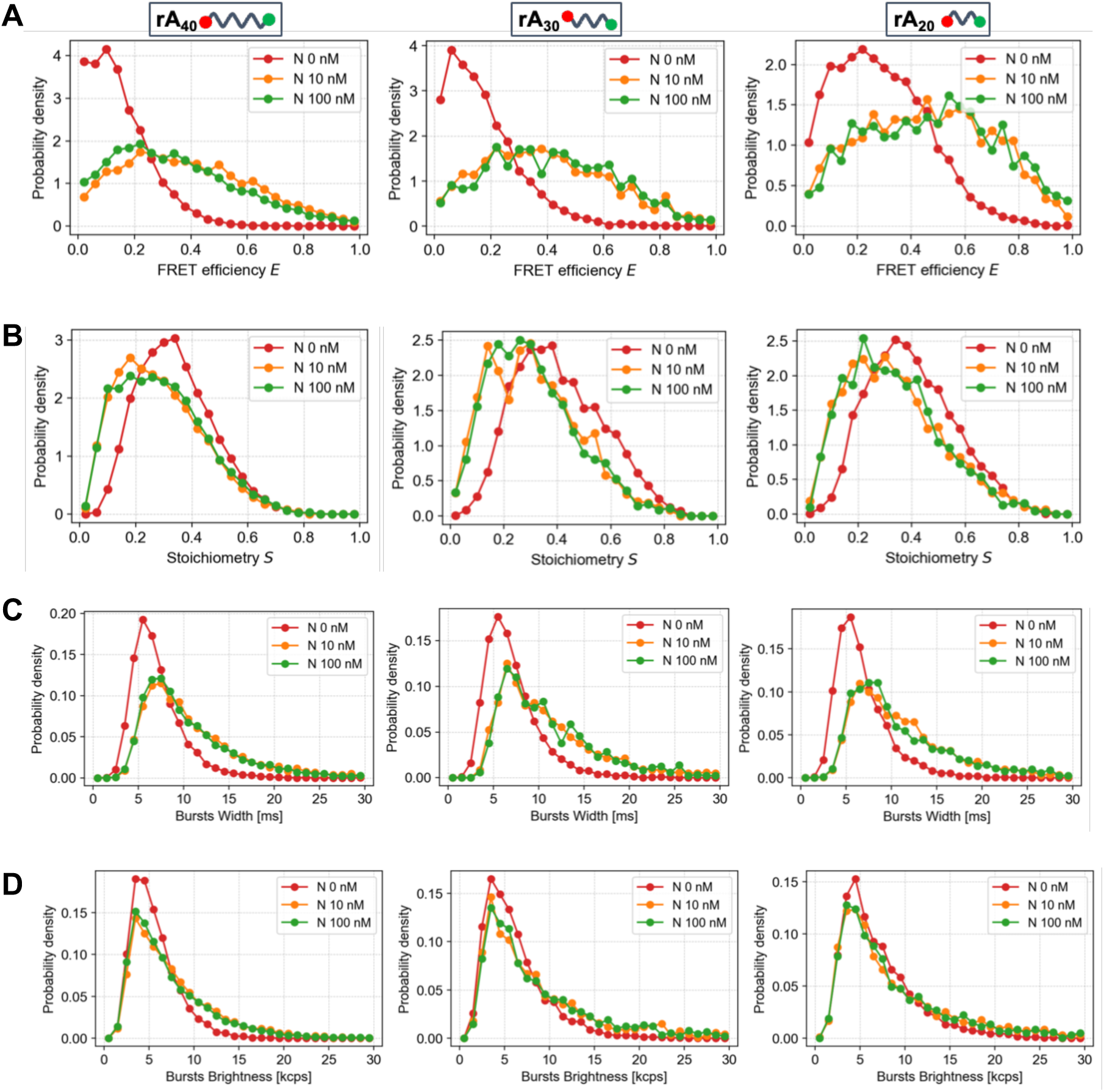
The sm-FRET data for the doubly-labeled single-stranded RNA fragments in the absence and presence of different concentrations of the N protein. A) Changes in the FRET efficiency distributions for 488-rA_40_-647 (left panel) at 100 pM, 488-rA_30_-647 (center panel) at 10 pM and 488-rA_20_-647 (right panel) at 25 pM in the absence (red), in the presence of 10 nM (orange) and 100 nM (green) of the N protein. B) Changes in the stoichiometry distributions. C) Changes in the burst width distributions. D) Changes in the burst brightness. The arrangement of the panels and the data colors in B, C and D are the same as those in A.

The association of the N protein to the RNA samples further caused changes in the brightness distribution of the burst signals (Figure 4D). To avoid a complication due to the quenching of Alexa488 fluorescence, we plotted the brightness for Alexa647 fluorescence obtained during the acceptor fluorophore excitation. In the absence of the N protein, the brightness was peaked at ∼4 kcps, and became ∼10 % of the peak value at ∼10 kcps. Since the sample is homogeneous in this condition, the distribution reflects the intensity distribution of the excitation laser and the detection efficiency distribution of the fluorescence photons in the observation volume. However, the addition of the N protein caused a decrease in the population having the brightness at ∼4 kcps and an increase in the population having the brightness at 10–15 kcps, the latter of which likely corresponds to the species having two or more Alexa647 in a single burst, consistent with the association of multiple RNA molecules to a single dimer of the N protein. The concentrations of the RNA samples for the sm-FRET measurements (25∼100 pM) were smaller than those used for the FCS measurements (0.5∼1 nM), likely explaining the relatively smaller contribution of the bursts having the higher brightness. Thus, the ALEX results for single-stranded RNAs were consistent with the suggestions from the CD and FCS measurements.

### Sm-FRET measurements of the doubly-labelled SL4, PS-SL1 and PS-SL2

Finally, we conducted the sm-FRET measurements for the RNA samples having the stem loop structures. The FRET efficiency distributions for 488-SL4-647, 488-PS-SL1-647 and 488-PS-SL2-647 were investigated in the presence of various concentrations of the N protein (Figure 5A). For all the samples in the absence of the N protein, the peak FRET efficiencies were higher than 0.8, indicating the proximity of Alexa488 and Alexa647 as expected for the stem loop structures. The binding of the N protein caused a slight decrease in the peak FRET efficiency and the broadening of the distribution. The donor-acceptor stoichiometry was 0.3 ∼ 0.4 in the absence of the N protein, but became lower and broader upon the addition of the N protein, supporting the quenching of the Alexa488 fluorescence (Figure 5B). The burst width distribution was peaked at ∼6 ms in the absence of N, but the addition of N caused the shift of the peak to 7–8 ms and the broadening of the distribution width, consistent with the increase of the hydrodynamic radius (Figure 5C). Finally, the distribution of the brightness of the acceptor fluorophore during the acceptor excitation was peaked at ∼5 kcps in the absence of the N protein. The addition of the N protein decreased the relative population of the bursts having the brightness of ∼5 kcps but increased the population having the brightness higher than 10 kcps (Figure 5D). Thus, except for the maintenance of the largely stem loop structures, the binding of the N protein resulted in the changes in the sm-FRET data sets for the stem loop samples very similar to those observed for single-stranded RNA samples.

**Figure 5.**
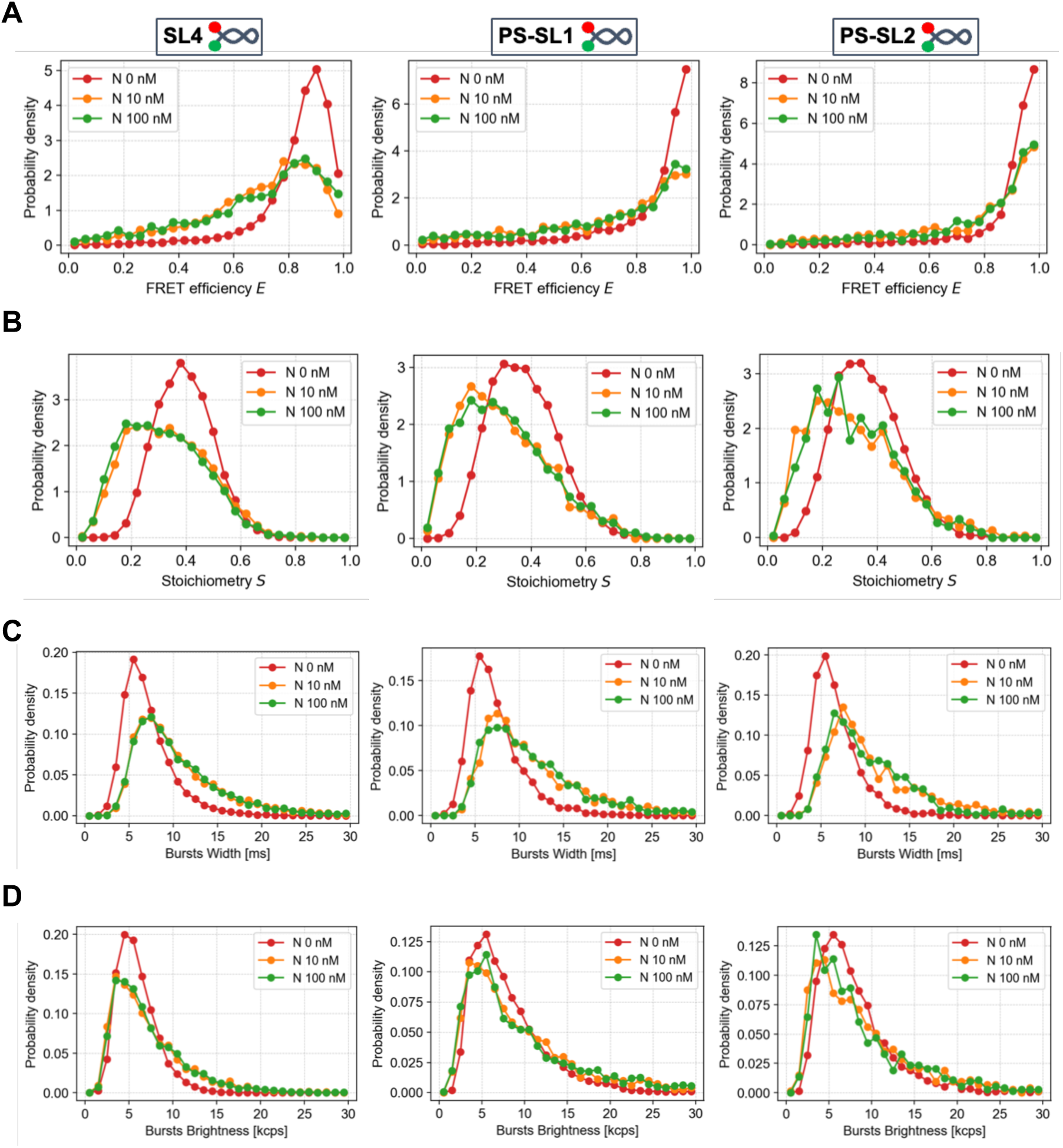
The sm-FRET data for the doubly-labeled stem-loop fragments of RNA in the absence and presence of different concentrations of the N protein. A) Changes in the FRET efficiency distributions for SL4 (left panel), PS-SL1 (center panel) and PS-SL2 (right panel) in the absence (red) and in the presence of 10 nM (orange) and 100 nM (green) of the N protein. All the RNA concentrations were 50 pM. B) Changes in the stoichiometry distributions. C) Changes in the burst width distributions. D) Changes in the burst brightness. The arrangement of the panels and the data colors in B, C and D are the same as those in A.

## DISCUSSION

### Association states of the single-stranded RNAs and the N protein

We will first discuss the association states and structural changes of the single stranded RNA sample, rA_40_. The CD spectrum for rA_40_ in the absence of the N protein showed the characteristic features corresponding to the π-stacked helical structure^45,46,47^. The single-molecule FRET measurements showed that 488-rA_40_-647 in the absence of the N protein possessed the peak FRET efficiency close to zero. Assuming a rod-like stricture, the FRET efficiency of 0.1 corresponds to the Alexa488-Alexa647 distance of ∼80 nm. The addition of the N protein reduced the CD ellipticities slightly, suggesting that the sample in the bound structure with the N protein keeps largely helical structure. The addition of the N protein increased the peak FRET efficiency and broadened the distribution for 488-rA_40_-647. The increase in the population of the bursts having the higher brightness suggested the binding of the multiple molecules of rA_40_ to a single dimer of the N protein. The FCS results showed that *R*_H_ of the N protein-rA_40_ complex is ∼8.7 nm. The FCS measurements based on the Alexa647 excitation showed that the addition of the N protein at 10 nM increased the *a* value ∼2 fold and reduced the *n*_R_ value to ∼0.5, suggesting that approximately two molecules of 488-rA_40_-647 was associated with a single dimer of the N protein. The changes in the *n*_R_ value was similarly detected for the FCS data excited at 482 nm and were consistent with the binding of multiple fragments of RNAs to a dimer of the N protein at 10 nM.

The systematic measurements of the FCS and sm-FRET data for rA_30_ and rA_20_ showed the same trend of results as those observed for rA_40_ except for the reduction of the affinity of rA_20_ towards the N protein. In the absence of the N protein, the sm-FRET peaks were commonly low, suggesting their extended helical structure. Upon the addition of the N protein, the *R*_H_ values became larger and the FRET efficiency peaks shifted to higher efficiency with significant broadening. The *n*_R_ values commonly became the smallest at the N protein concentration of 10 nM, suggesting the association of the multiple molecules of the RNAs to a single dimer of the N protein. Based on these results, we depicted a plausible model for the binding of the single-stranded RNA to the N protein (Figure 6A). The binding of the N protein and the RNA samples near the N protein concentration at 10 nM might be in line with the dimerization of the N protein, as the dissociation of a N protein dimer to monomers was reported to occur at a few nM concentration^51^.

**Figure 6.**
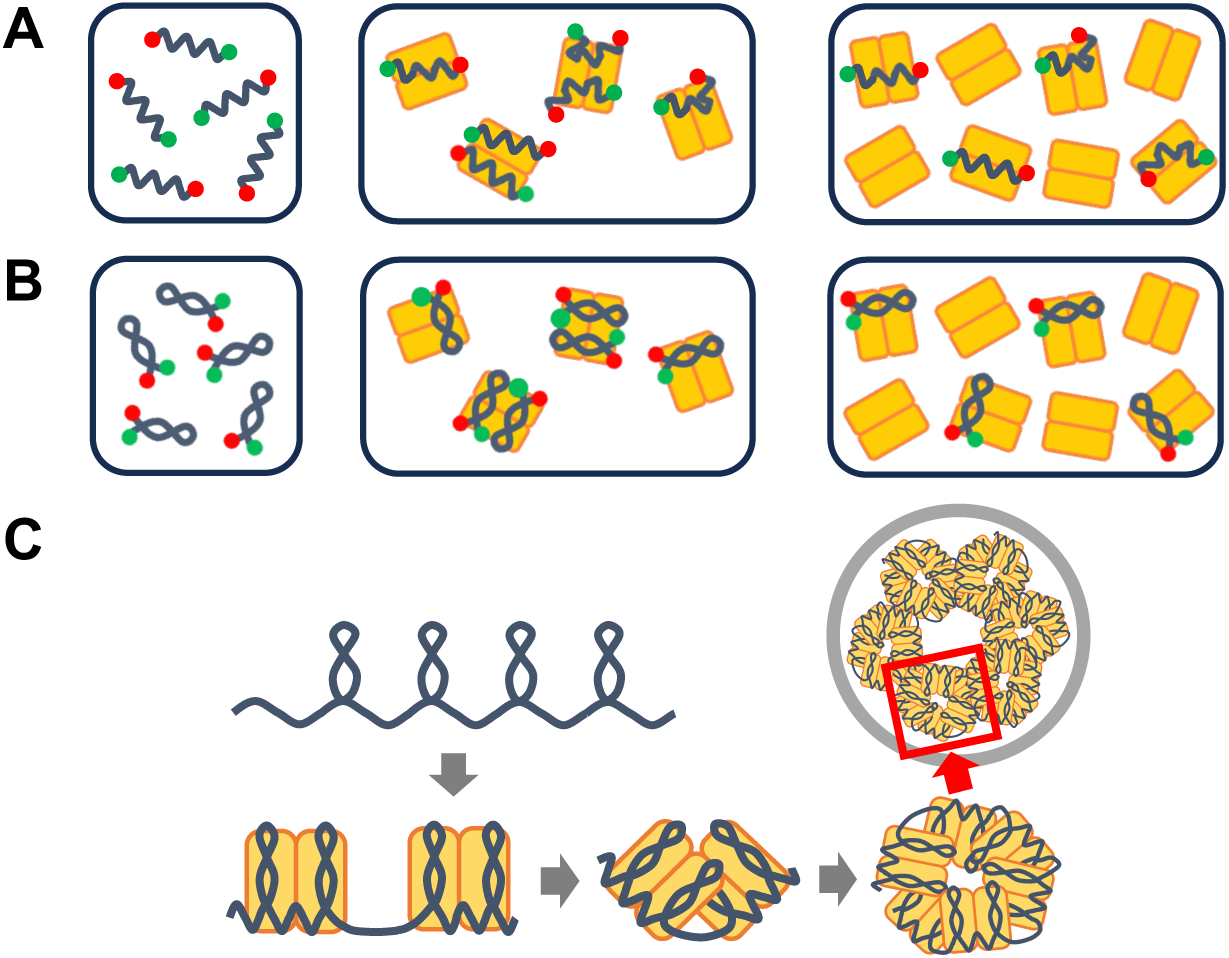
Folding model of the RNP granule formation. A) Single stranded RNAs associate with the N protein in the mostly helical structure. Multiple fragments of single stranded RNA bind to a single dimer of the N protein. B) Stem loop RNAs associate with the N protein without melting the stem loops. More than two molecules of RNA bind to a single dimer of the N protein. C) Folding model of the RNP granule formation. The stem loops of gRNA formed autonomously are bridged by the N protein dimer, facilitating the long-range base pairing and the folding of gRNA granules.

### Association state of the double-stranded RNAs and the N protein

We will next discuss the properties of the double-stranded RNAs upon the association with the N protein. The sm-FRET results for 488-SL4-647 in the absence and presence of the N protein demonstrated the peak efficiencies close to one, showing the proximity of the 3’ and 5’ ends consistent with the stem loop structure. The CD spectra of SL4 in the absence and presence of the N protein overlapped perfectly, demonstrating that the stem loop structure of SL4 does not change upon the binding with the N protein. The broadening of sm-FRET distribution of 488-SL4-647 upon the binding of the N protein might be explained by factors other than the melting of the stem loop such as the restricted rotation of the fluorophores. The FCS measurements at the Alexa647 excitation showed that the addition of the N protein increased the SL4 brightness for more than ∼5 fold. While the value contains a rather large error, it suggested the association of more than two molecules of SL4 to the N protein dimer. Thus, while the binding stoichiometry might be different for rA_40_ and SL4, the current results for SL4 showed similar trend of results for the single-stranded and stem loop structures of RNA. A possible model for the binding of SL4 to the N protein was depicted in Figure 6B. We further examined the two stem-loop structures, PS-SL1 and PS-SL2, which were proposed as possible candidates of the packaging signals^44^. The packaging signal is a hypothetical sequence of gRNA that might interact specifically with the N protein and trigger the selective packaging of a single gRNA inside of the virus by associating with the M protein^31^. The single-molecule FRET results for the two PS samples were almost identical to those for SL4, showing that the association pattern of these stem loops to the N protein is similar to that of SL4.

### Folding model of the RNP granule formation induced by the association of gRNA and the N protein

Based on the current investigation, we propose the folding model of the RNP formation. The cryo-electron tomography showed that 35∼40 granules of RNP having a diameter of ∼15 nm were filled inside of the SARS-CoV-2^4,7,8^. The current investigation indicated that the N protein dimer possesses the binding sites for more than two fragments of RNA and that the stem-loop fragments possess a higher affinity to the N protein relative to the short single-stranded fragments. Thus, we can assume that the N protein dimer can contract the long gRNA by binding to its two or more stem-loop structures. It was previously shown that gRNA inside of the virus possesses a large number of stem loops^13,14^. As we demonstrated that the autonomous formation of the three stem loop structures, all the stem loops should be formed in the full-length gRNA. The N protein can promote the contraction of the multiple stem loops of gRNA by bridging between them and might lead to the formation of the long-range base pairing having the separation of several hundreds of bases reported in gRNA^13^. We suggest that the association of the N protein dimer glues the stem loops and triggers the folding of the RNP granules (Figure 6C).

## Conclusion

In this study, CD spectroscopy, FCS and sm-FRET spectroscopy were used to investigate the association states and structural changes of several short fragments of RNA upon the binding with the SARS-CoV-2 N protein. The polyadenylate chains with different lengths, rA_40_, rA_30_ and rA_20_ and the fragments selected from the stem loop regions of gRNA, SL4, PS-SL1 and PS-SL2 were labeled with fluorophores for the FCS and FRET measurements. The CD spectroscopic results showed that the binding of the N protein only slightly reduced the single-stranded helical content of rA_40_, and did not modulate the stem loop structure of SL4. The FCS results demonstrated that approximately two molecules of rA_40_ associate with a single dimer of the N protein at the N protein and RNA concentrations of 10 nM and 1 nM, respectively. In the case for SL4, more than two molecules of SL4 binds to a single dimer of the N protein at the same concentrations. The sm-FRET results further supported the binding of multiple RNA molecules to a single dimer of the N protein. Based on the current results, the folding model of the RNP granule formation was proposed, in which the N protein triggers the folding of gRNA to form the granular RNP structures.

## Notes

### Competing Interest Statement

The authors have declared no competing interest.

