## Supporting Information for "A single dimer of the SARS-CoV-2 N protein can associate with multiple fragments of single-stranded and stem-loop RNAs"

### Supporting Texts

***Sample preparation and reagents.*** Fluorescently-labeled RNA oligos were purchased from Ajinomoto Bio-Pharma Services (Osaka, Japan). All chemicals used were commercially available.

***Construction of the expression system for the N protein of SARS-CoV-2 and its purification.*** We constructed the expression system for the N protein whose N-terminus is tagged with 10 histidines (His10). The gene for His10-N was cloned into the pET28a vector by Eurofins Genomics. The vector was transformed into *E. coli*, BL21(DE3). The cells were grown in 10 mL Luria Broth (LB) media containing 50 μg/mL kanamycin at 37°C for 17 h, and transferred into 1 L LB media containing 50 μg/mL kanamycin and were grown at 37°C until the optical density at 600 nm reached ~0.6. The cells were further cultured for 4 h after adding isopropyl β-D-1-thiogalactopyranoside to the final concentration of 1 mM, and were harvested by centrifugation at 7000 rpm at 4°C for 5 min. The pellet was frozen and stored at –80°C. The cell pellet was suspended in 40 mL of the sonication buffer containing 20 mM tris(hydroxymethyl)aminomethane (Tris)-HCl at pH 8.0, 200 mM NaCl, and 12.5 μM ribonuclease (RNase) A (312-01931, Nippon Gene). The cells were lysed by 5-min sonication three times by using the ultrasonic disruptor (UD-201, Tomy Seiko). The solution was centrifuged at 12000 rpm at 4°C for 30 min. The pellet was washed three times by resuspending it with 40 mL of the first suspension buffer containing 20 mM Tris-HCl at pH 8.0, 200 mM NaCl, 12.5 μM RNase A and 0.3% Tween-20, and by centrifuging the suspension at 12000 rpm at 20°C for 30 min. The pellet was resuspended in the second suspension buffer containing 20 mM Tris-HCl at pH 8.0 and 200 mM NaCl, and centrifuged at 12000 rpm at 4°C for 30 min. The pellet was resuspended overnight with 10 mL of the urea buffer containing 20 mM Tris-HCl at pH 7.5, 1 M NaCl and 6 M urea. The N protein was purified in the unfolded state by using the NGC system (Bio-Rad). The resuspended solution was passed through a Ni-NTA affinity column (Histrap$^{TM}$ FF crude 1 mL, Cytiva) and washed with the wash buffer containing 20 mM Tris-

HCl at pH 7.5, 1 M NaCl, 6 M urea and 50 mM imidazole. The N protein was eluted with the elution buffer containing 20 mM Tris-HCl at pH 7.5, 1 M NaCl, 6 M urea and 500 mM imidazole. To remove urea from the solution, the N protein solution was dialyzed overnight in the storage buffer containing 20 mM Tris-HCl at pH 7.5, 1 M NaCl and 2 mM $MgCl_2$ by using the dialysis cassette (7000 cut off, 87727, Thermo Fisher Scientific). The concentration of the N protein was estimated by measuring the absorbance at 280 nm using Nanodrop (Thermo Fisher Scientific), and the molar extinction coefficient ($\varepsilon = 43890$ L·mol$^{-1}$·cm$^{-1}$) estimated from the numbers of the aromatic amino acids. The N protein was divided into small aliquots and flash frozen in liquid nitrogen and kept at –80°C.

***CD spectroscopy.*** A CD spectrum of the N protein at 5.2 μM in the measurement buffer containing 20 mM Tris-HCl at pH 7.5, 150 mM NaCl and 2 mM $MgCl_2$ was measured with a spectropolarimeter (J-720, Jasco). The CD spectra of 5 μM SL4 and 2 μM $rA_{40}$ RNA in the presence of 0 and 2 μM of the N protein were measured in the same condition. The spectra in the presence of 5 μM of the N protein were measured using the buffer containing 20 mM Tris-HCl at pH 7.5, 319 mM NaCl and 2 mM $MgCl_2$. All measurements were performed with a cell having 1 mm pathlength at room temperature.

***Detection systems for the FCS measurements.*** For the FCS measurements based on the 484-nm excitation, we used a hand-made FCS spectrometer described previously[1]. Briefly, a 484-nm laser light (LDS1003, Precise Gauges) was focused to the sample solution by a water immersion objective (UPlanSApo 60x, Olympus). The fluorescence photons were collected by the same objective, passed through the first dichroic mirror and focused to a 50-μm pinhole (P50H, Thorlabs). The photons were further passed through the second dichroic mirror (67-083, Edmund) and the bandpass filter (FBH520-40, Thorlabs), and were detected by HPD (H13223-40, Hamamatsu Photonics). The HPD outputs were amplified (C10778, Hamamatsu Photonics) and recorded by a time correlated single photon counting

(TCSPC) module in the time-tag mode (TimeHarp 260 NANO, PicoQuant) controlled by a software (QuCoa, PicoQuant). The time-tagged photon data were converted into the photon intensity correlations using a software (SymPhoTime64, PicoQuant). The reproducibility of fluorescence intensity measurements of the system estimated as the standard deviation of the multiple brightness measurements of Alexa488 sampled from the same stock solution in the same day, same laser power and same optical alignment was 2.4 % of the averaged brightness. For the FCS measurements based on the 642 nm excitation, we used the system developed previously for the ALEX measurements of the sm-FRET spectroscopy with some modifications as explained below[2]. The fluorescence intensity signals were converted into the correlations by using a home-made software based on LabView[3,4]. The reproducibility of fluorescence intensity measurements of the system estimated as the standard deviation of the multiple brightness measurements of Alexa647 sampled independently from the same stock solution in the same day, same laser power and same optical alignment was 2.6 % of the averaged brightness.

***FCS measurements.*** The measurement buffer contained 20 mM Tris-HCl at pH 7.5, 150 mM NaCl and 2 mM $MgCl_2$. As the reference samples, ~1 nM solutions of rhodamine 110 in pure water and Alexa488 in the measurement buffer were prepared. The concentration of the labeled RNA was adjusted at ~1 nM and that of N protein was between 0 nM and 1000 nM. The inner surface of glass base dishes (3971-035, AGC Techno Glass) was coated with 2-methacryroyloxyethylphosphorylcholine (MPC) polymer (Lipidure CM5206, NOF) by rinsing the surface by 0.5% (w/v) solution of MPC in 99.5% ethanol followed by drying. 300 μL of the sample solution was placed on the coated dish, and was covered with a lid. The FCS measurements were performed in a room adjusted at 22°C. The excitation laser power was measured near the focus point of the objective with a power meter (PM400, Thorlabs), and was adjusted to 50 μW. The data acquisition time for one measurement was 10 min. All measurements were conducted in a room temperature. For the FCS measurements based on the 642 nm excitation, we used

~1 nM solution of Alexa647 in the measurement buffer as the reference sample. To obtain the data in the same measurement conditions, we optimized and tightened the optical configuration using rhodamine or free Alexa647 solutions and conducted a series of measurements of one RNA sample at the different concentrations of the N protein in the same day without changing the optical configuration, the laser setting and the cell stages.

***Fitting of the correlation functions and FCS data analysis.*** Preliminary FCS measurements of the RNA samples labeled with Alexa488 at the 484-nm excitation showed that the time constant for the triplet state accumulation was a few microseconds at the laser power of 50 μW, and that the triplet state in the correlogram in the time domain longer than 30 μs was negligible. Thus, to minimize the number of the fitting parameters, we conducted the FCS measurements at the excitation of 50 μW and fitted the correlogram in the time domain from 30 μs to 100 ms without considering the triplet state contribution. In the case for the FCS measurements for the samples labeled with Alexa647 at the 642-nm excitation, the correlograms further contained another phase ascribed to the cis-trans isomerization of the fluorophore occurring with a time constant of 1~20 μs[5]. To completely eliminate the phase ascribed to the isomerization, we fitted the correlograms in the time domain from 100 μs to 100 ms.

As we explained in the main text, the correlation functions, $G(\tau)$, were fit with eq. S1, assuming the presence of two diffusing components having different diffusivity and brightness, one is the free Alexa488 diffusing quickly with a constant brightness and the other is the RNA samples labeled with Alexa488 diffusing slowly with a variable brightness (eq. S1):

$$G(\tau) = \frac{b^2}{\langle I \rangle^2} \cdot \left\{ \left( \frac{\langle I \rangle}{b} - a \cdot n_R \right) \cdot g_A(\tau) + a^2 \cdot n_R \cdot g_R(\tau) \right\} + 1 \qquad \text{eq. S1,}$$

where $b$, $<I>$, $a$, $n_R$ are the brightness of free Alexa488, the averaged fluorescence intensity during the FCS measurement, the ratio of the brightness of the labeled RNA relative to that of the free fluorophore and the number of the labeled RNA in the observation volume, respectively. $<I>$ can be expressed as follows:

$$\langle I \rangle = b \cdot (n_A + a \cdot n_R) \qquad \text{eq. S2,}$$

where $n_A$ is the number of the free fluorophore in the observation volume. Eq. S1 can be derived from the standard equation for the two-component system (eq. 1 in the main text) by using eq. S2[6,7]. Eq. S2 was also used to estimate $n_A$ from $<I>$ and other fitted parameters. $g_A(\tau)$ and $g_R(\tau)$ are the normalized correlations for the translational diffusion of the free fluorophore and the labeled RNA, respectively, and are expressed as follows:

$$g_{A\ or\ R}(\tau) = \frac{1}{1 + \tau/\tau_{A\ or\ R}} \cdot \frac{1}{\sqrt{1 + s^{-2} \cdot \tau/\tau_{A\ or\ R}}} + 1 \qquad \text{eq. S3,}$$

where $s$ is the ratio of the axial radius to the radial radius of the observation volume, and $\tau_A$ and $\tau_R$ are the translational correlation times for the free fluorophore and the labeled RNA, respectively.

We conducted the standard least-squared fitting using a custom-made Python scripts implementing eqs. S1 ~ S3, utilizing the curve_fit function from scipy.optimize (SciPy v 1.10.1). The $s$ factor was fixed to 8.40, 7.50, 8.00 and 7.10 in the fitting of the autocorrelations obtained for the Alexa488-labeled SL4, $rA_{40}$, $rA_{30}$ and $rA_{20}$, respectively. The $s$ factor was fixed to 11.0 and 11.2 in the fitting of the autocorrelations excited at 642 nm for SL4 and $rA_{40}$, respectively. We conducted the FCS measurements of free Alexa488 and obtained that the $\tau_A$ value was 79 μs. In the case for free Alexa647,

$\tau_A$ was either 198 or 195 μs. We confirmed that the $\tau_A$ values of free Alexa488 and free Alexa647 did not change in the absence and presence of the N protein at 100 nM, demonstrating the absence of the direct interaction between the free fluorophores and the N protein. We assumed that the brightness of free Alexa is the same as that of the RNA labeled Alexa in the absence of the N protein, that is, *a* is 1, and fitted the correlation and fluorescence intensity data in the absence of the N protein using eq. S1, and obtained *b*, $n_R$ and $\tau_R$. For the data in the presence of the N protein, the *b* parameter obtained in the absence of the N protein was fixed to obtain *a*, $n_R$ and $\tau_R$. The reduction of *a* upon the addition of the N protein might imply the quenching of the Alexa fluorescence labeled to RNA and the increase of *a* the association of multiple RNA labeled with fluorophore to a single particle of the N protein.

Using the $\tau_R$ value estimated by the fitting, we calculated the hydrodynamic radius, $R_H$, of the labeled RNA by converting $\tau_R$ to a diffusion coefficient, $D_R$, using eq. S4 and by converting $D_R$ to $R_H$ using eq. S5:

$$D_R = \frac{{\omega_0}^2}{4 \cdot \tau_R} \qquad \text{eq. S4,}$$

$$R_\mathrm{H} = \frac{k_B \cdot T}{6 \cdot \pi \cdot \eta \cdot D_R} \qquad \text{eq. S5,}$$

where $k_B$ is the Boltzmann constant ($1.38 \cdot 10^{-23}$ J/K), *T* is the absolute temperature set at 293 K and $\eta$ is the viscosity ($9.58 \cdot 10^{-4}$ Pa·s), and $\omega_0$ is the short radius of the focus area. The $\omega_0$ value was determined using the reference FCS data for free fluorophores using their reported diffusion coefficients, 435 $\mu m^2/s$ for Alexa488 and 330 $\mu m^2/s$ for Alexa647[8,9].

The reproducibility of the fluorescence intensity measurements of the FCS system for the 484-nm excitation estimated as the standard deviation of the brightness of Alexa488 was 2.4 % of the averaged brightness. Similarly, the standard deviation of the brightness of Alexa647 determined by the FCS system for the 642-nm excitation was 2.6 % of the averaged brightness. Thus, the systems could reasonably reproduce the fluorescence intensities if the multiple measurements were conducted in the same day and in the same optical set up. However, the fluorescence intensity might still change due to the sample concentration changes caused by sampling errors and/or by adsorption to cell surfaces. While we conducted the sample preparations carefully and used glass base dishes freshly coated by an anti-adsorption MPC polymer, the sample concentration might still change. To estimate the possible errors caused by the sample concentration fluctuation, we assumed ± 10 % errors in $\langle I \rangle$ and estimated the changes of the fitting parameters and plotted them as the possible error ranges.

***Apparatuses for the ALEX measurements.*** We performed ALEX measurements by using a home-made confocal microscope reported previously with some modifications[2]. The system is equipped with two lasers, one for the excitation of donor at 488 nm (OBIS 488 nm LX 100 mW, 1236444, Coherent) and the other for the acceptor at 642 nm (OBIS 640 nm LX 75 mW, 1236445, Coherent) assembled in a laser box (OBIS LX Laser Box, 1228877, Coherent). The outputs of the two lasers were combined by using a combiner (OBIS Galaxy, 1363484, Coherent), collimated by an achromatic collimator (C80FC-A, Thorlabs) and passed through an iris (SM2D25, Thorlabs). The excitation beam was reflected by a wedge plate (BSF10-A, Thorlabs), introduced into a water immersion objective (NA 1.2, CFI PlanApo 60XC WI, Nikon), and focused into the sample placed on the glass-based dish (IWAKI). Fluorescence photons were collected by the same objective, passed through the wedge plate and a dual bandpass filter (ZET488/640m, Chroma Technology), and were separated into donor and acceptor fluorescence by a dichroic mirror (593 nm cut-on wavelength, #67-083, Edmund Optics). The separated photons were passed through bandpass filters (for donor fluorescence, FBH520-40, Thorlabs, and for acceptor fluorescence, FF01-676/29, Semrock) and were imaged with two achromatic lenses (AC254-200-A-ML, Thorlabs) onto multimode optical fibers with a core diameter of 100 μm. The fibers were coupled to two single-photon avalanche diodes (SPADs, SPCM AQRH-14-FC, Excelitas). Signals from SPADs were recorded as photon arrival times by using a counter (PCIe-6612, National Instruments) operated by home-built LabVIEW software (National Instruments). For all the ALEX measurements, the excitation lasers were modulated with square waves at a frequency of 50 kHz and a duty ratio of 0.5. The excitation laser power was measured near the focus point of the objective with a power meter (Thorlabs). The donor excitation laser power was adjusted to 20 μW for $rA_{40}$, SL4, PS-SL1 and PS-SL2, and to 60 μW for $rA_{20}$ and $rA_{30}$, respectively. The acceptor excitation laser power was 10 μW for $rA_{40}$, SL4, PS-SL1 and PS-SL2, and 30 μW for $rA_{20}$ and $rA_{30}$, respectively.

***ALEX measurements.*** The concentration of $rA_{40}$, $rA_{30}$, and $rA_{20}$ was 100, 10, and 25 pM, respectively, and that of SL4, PS-SL1, and PS-SL2 was 50 pM. The N protein concentration was 0–100 nM. The measurement buffer contained 20 mM Tris-HCl at pH 7.5, 150 mM NaCl and 2 mM $MgCl_2$. The solutions of the double-labeled RNA samples and the N protein were mixed in the measurement buffer immediately before the ALEX measurements. 400 μL of the solution was placed on the glass-based dish, or 200 μL of the solution was placed on the coverslip (NO.1 SHT, Matsunami Glass Ind., Ltd.) coated with MPC. For the MPC coating, we dropped the 0.5% (w/v) solution of MPC in 99.5% ethanol onto the coverslip and kept it till the MPC solution dried. The ALEX measurements were performed in a room adjusted at 22°C. To prevent the adsorption of RNA and N protein, the surfaces of the glass-based dishes were coated with MPC polymer as described above. The total data acquisition time was 15−30 min. All measurements were conducted in a room temperature.

***ALEX data analysis.*** The arrival times of all the photons stored in the photon-HDF5 file format[10] were analyzed by using a software, FRETBursts[11]. First, the local background rates for each of the three photons, the photons detected in the donor channel during the donor excitation, the photons detected in the acceptor channel during the donor excitation and the photons detected in the acceptor channel during the acceptor excitation were estimated at every 30 s window, whose typical values were ~0.3, ~0.2 and ~0.4 kilocounts per second (kcps), respectively. Second, the fluorescence bursts were searched for the three channels by selecting the time regions where the local count rate calculated using arrival times of 10 consecutive photons exceeded the count rate thresholds that were set to the averaged background rate plus 4 kcps. This threshold was necessary for the data reproducibility. For each of the selected bursts, there types of the photon numbers were estimated after the subtraction of the local background rates, which include the number of the donor photons during the donor excitation ($F_{\mathrm{D_{ex}}}^{\mathrm{D_{em}}}$), the number of the acceptor photons during the donor excitation ($F_{\mathrm{D_{ex}}}^{\mathrm{A_{em}}}$) and the number of the acceptor photons during the acceptor excitation ($F_{\mathrm{A_{ex}}}^{\mathrm{A_{em}}}$). Third, the bursts were further selected by using the threshold of 15 for the sum of donor and acceptor photons obtained during the donor excitation, $F_{\mathrm{D_{ex}}}^{\mathrm{D_{em}}} + F_{\mathrm{D_{ex}}}^{\mathrm{A_{em}}}$, to eliminate the acceptor-only species. Fourth, another selection was performed by setting the threshold of 15 for $F_{\mathrm{A_{ex}}}^{\mathrm{A_{em}}}$ to eliminate the donor-only bursts. The FRET efficiency (*E*) and stoichiometry (*S*) for each burst were calculated using eqs. S6 and S7, respectively:

$$E = \frac{F_{\mathrm{D_{ex}}}^{\mathrm{A_{em}}}}{\gamma \cdot F_{\mathrm{D_{ex}}}^{\mathrm{D_{em}}} + F_{\mathrm{D_{ex}}}^{\mathrm{A_{em}}}} \qquad \text{eq. S6,}$$

$$S = \frac{\gamma \cdot F_{\mathrm{D_{ex}}}^{\mathrm{D_{em}}} + F_{\mathrm{D_{ex}}}^{\mathrm{A_{em}}}}{\gamma \cdot F_{\mathrm{D_{ex}}}^{\mathrm{D_{em}}} + F_{\mathrm{D_{ex}}}^{\mathrm{A_{em}}} + F_{\mathrm{A_{ex}}}^{\mathrm{A_{em}}}} \qquad \text{eq. S7,}$$

where $\gamma$ represent the ratio of the florescence detection efficiencies for the donor and acceptor channels and was set to a previously estimated value of 0.48[2]. The stoichiometry $S$ depends on the relative intensities of the two lasers used to excite the donor and acceptor. The peak stoichiometry values (0.3~0.4) observed for the RNA samples in the absence of the N protein correspond to the samples containing the donor and acceptor at the 1:1 ratio.

### Supporting Figures

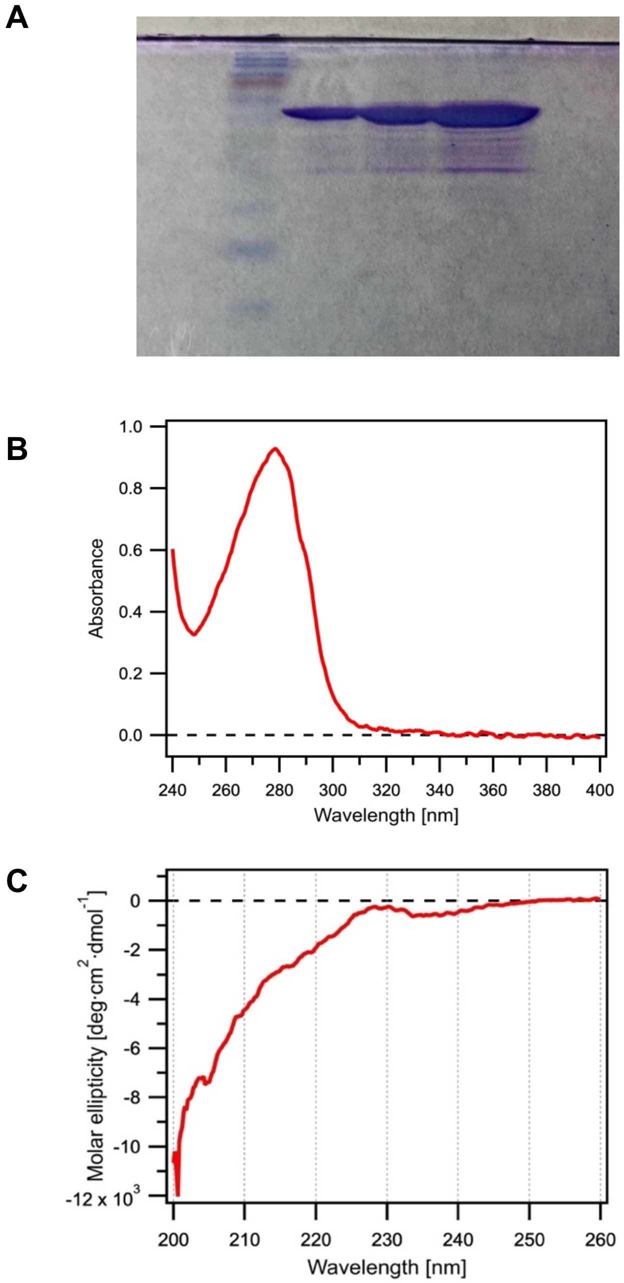


**Supporting Figure S1. Purity of the N protein preparation.** A) The SDS-PAGE pattern for the final purified sample of the N protein. The impurity components were detectable only in the heavily lorded lane. The N protein appeared between two markers corresponding to 50 and 60 kDa. B) The optical absorption spectrum of the purified N protein in the UV and visible wavelength regions. C) The circular dichroism spectrum of the purified N protein at 5.2 μM in the far UV region.

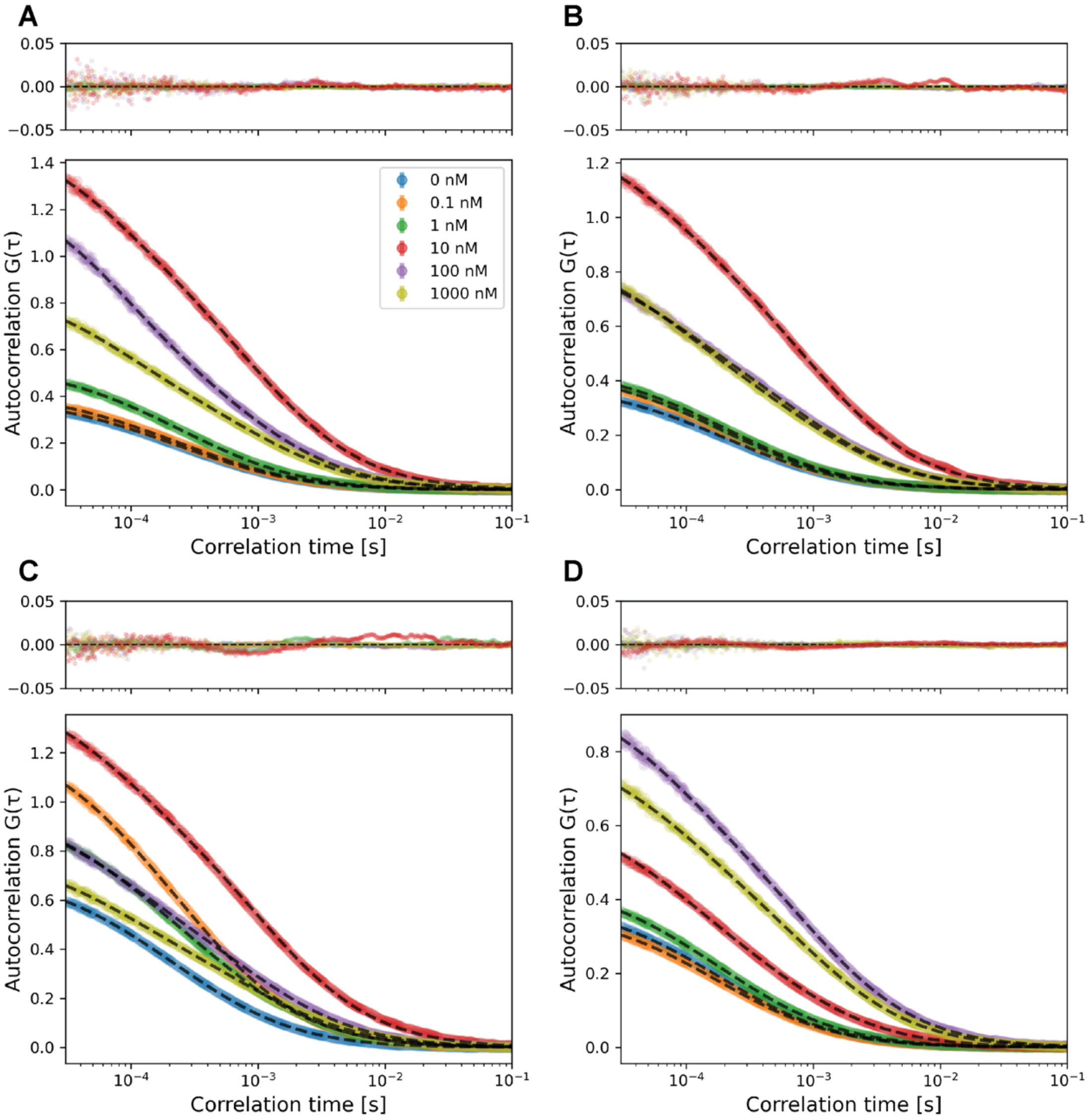


**Supporting Figure S2. The FCS correlograms and the fitted curves for 488-rA$_{40}$, 488-rA$_{30}$, 488-rA$_{20}$ and 488-SL4 at different concentrations of the N protein.** The FCS data were obtained at the 484-nm excitation, and analyzed assuming two diffusing components, the free Alexa488 diffusing quickly and the labeled RNA diffusing slowly. For all the panels, the data obtained at 0 nM, 0.1 nM, 1 nM, 100 nM and 1000 nM of the N protein were presented in cyan, orange, green, red, purple and olive, respectively. The best fitting curves were presented in black broken line. Panels A, B, C and D are the data obtained for 488-rA$_{40}$, 488-rA$_{30}$, 488-rA$_{20}$ and 488-SL4, respectively. The residuals of the fitting were presented in the upper sub panels.

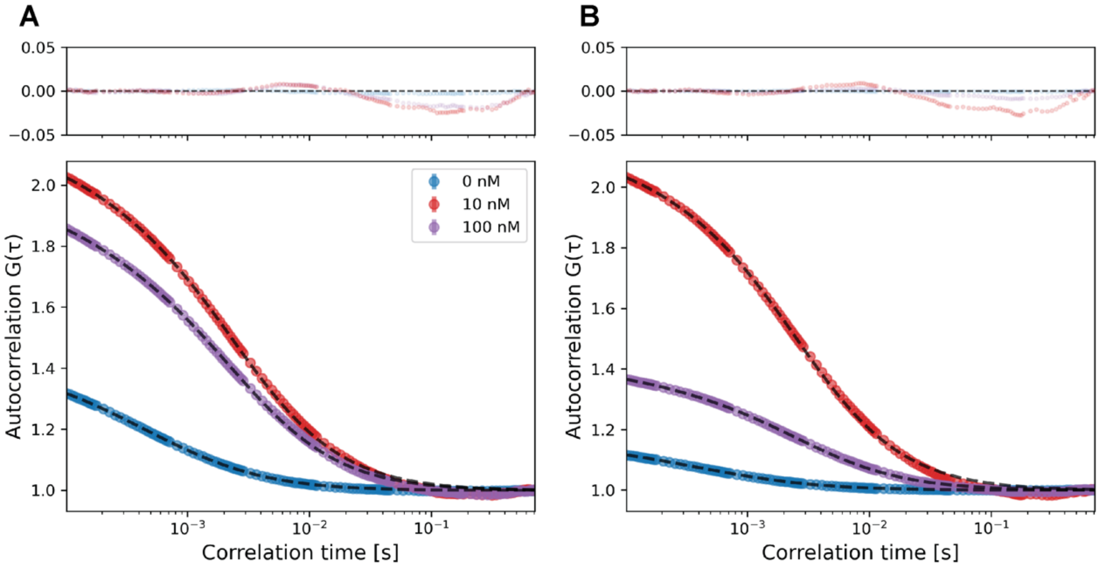


**Supporting Figure S3. The FCS correlograms and the fitted curves for 488-rA$_{40}$-647 and 488-SL4-647 at different concentrations of the N protein.** The FCS data were obtained at the 642-nm excitation, and analyzed assuming two diffusing components, the free Alexa647 and the labeled RNA. For all the panels, the data obtained at 0 nM, 10 nM and 100 nM of the N protein were presented in cyan, red and purple, respectively. The best fitting curves were presented in black broken line. Panels A and B are the data obtained for 488-rA$_{40}$-647 and 488-SL4-647, respectively. The residuals of the fitting were presented in the upper sub panels.

A

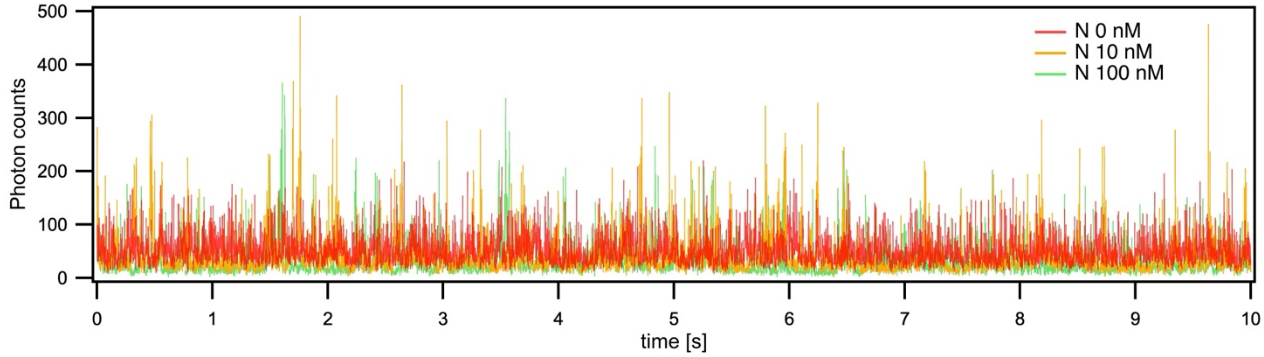


B

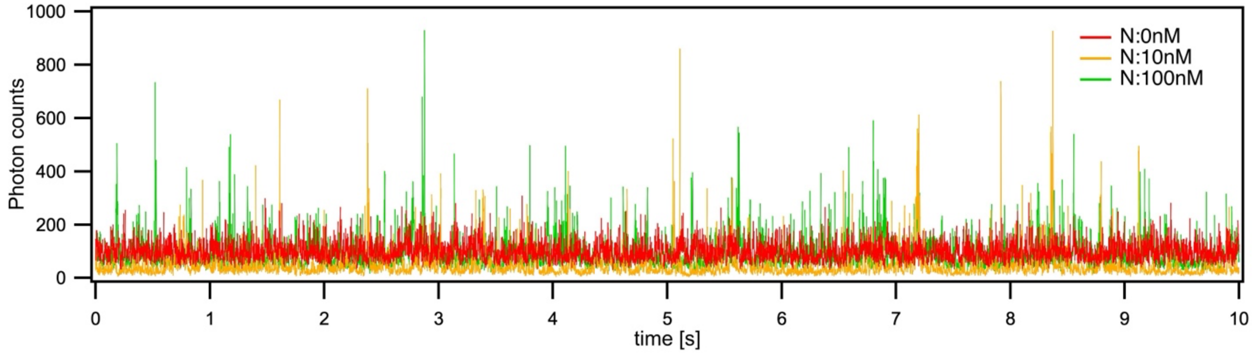


**Supporting Figure S4. The fluorescence intensity fluctuations for 488-rA$_{40}$-647 (A) and 488-SL4-647 (B).** Part of the raw fluctuation data were presented that were obtained in the FCS measurements excited at 642 nm at 0 nM (red), 10 nM (orange) and 100 nM (green) of the N protein.

**Supporting Table S1A. The parameters used in and obtained by the fitting analysis of the FCS data for 488-rA$_{40}$ excited at 484 nm**[1)]

| Concentration[2)] | 0 nM | 0.1 nM | 1 nM | 10 nM | 100 nM | 1000 nM |
|---|---|---|---|---|---|---|
| $<I>$[3)] | 39.0 ± 3.9 photons/ms | 36.2 ± 3.6 photons/ms | 26.6 ± 2.7 photons/ms | 8.69 ± 0.87 photons/ms | 7.10 ± 0.71 photons/ms | 10.5 ± 1.1 photons/ms |
| $b$[4)] | 15.3 ± 1.5 photons/ms | 15.3 photons/ms | 15.3 photons/ms | 15.3 photons/ms | 15.3 photons/ms | 15.3 photons/ms |
| $\tau_R$[5)] | 418 ± 1.9 μs | 460 ± 2.1 μs | 419 ± 2.5 μs | 1150 ± 4.2 μs | 1060 ± 7.9 μs | 1160 ± 5.5 μs |
| $n_R$[6)] | 1.55 ± 0.01 | 1.40 ± 0.20 | 1.28 ± 0.14 | 0.45 ± 0.05 | 0.78 ± 0.10 | 1.15 ± 0.12 |
| $a$[7)] | 1.00 | 0.98 ± 0.17 | 0.89 ± 0.14 | 0.79 ± 0.13 | 0.36 ± 0.06 | 0.40 ± 0.06 |
| $\tau_A$[8)] | 79 μs | 79 μs | 79 μs | 79 μs | 79 μs | 79 μs |
| $n_A$[9)] | 1.00 ± 0.36 | 0.99 ± 0.39 | 0.60 ± 0.27 | 0.21 ± 0.09 | 0.19 ± 0.08 | 0.23 ± 0.11 |
| $s$[10)] | 7.5 | 7.5 | 7.5 | 7.5 | 7.5 | 7.5 |
| $R_H$[11)] | 3.23 ± 0.28 nm | 3.56 ± 0.30 nm | 3.25 ± 0.28 nm | 8.87 ± 0.76 nm | 8.23 ± 0.70 nm | 8.95 ± 0.76 nm |

[1)]The FCS data obtained for 488-rA$_{40}$ excited at 484 nm was analyzed based on eqs. S1 ~ S3. The definitions of the parameters were explained in Supporting Texts. The FCS data and fitted curves were presented in Supporting Figure S2A. The errors of the parameters were estimated by the method explained in Supporting Texts. [2)]The N protein concentration. [3)]The averaged fluorescence intensity during the FCS measurements used for the fitting. The error ranges were roughly estimated as 10 % of the measured fluorescence intensity. [4)]The brightness of the free Alexa. The value estimated for the data in the absence of the N protein assuming $a = 1$ was used as the fixed fitting parameter for the data in the presence of the N protein. [5)]The translational diffusion time for the labeled RNA obtained by the fitting. [6)]The number of the labeled RNA in the observation volume obtained by the fitting. [7)]The ratio of the brightness of the labeled RNA relative to that of the free Alexa. The value was fixed to 1 for the data in the absence of the N protein to estimate $b$, and set as a fitting parameter for the data in the presence of the N protein. [8)]The translational diffusion time for the free Alexa. The value was fixed to the value determined separately. [9)]The number of the free Alexa in the observation volume estimated using eq. S2 and other fitted parameters. [10)]The ratio of the axial radius to the radial radius of the observation volume. The value was fixed to the value determined separately. [11)]The hydrodynamic radius of the labeled RNA estimated from $\tau_R$ using eqs. S4 and S5.

**Supporting Table S1B. The parameters used in and obtained by the fitting analysis of the FCS data for 488-rA$_{30}$ excited at 484 nm[1)]**

| Concentration[2)] | 0 nM | 0.1 nM | 1 nM | 10 nM | 100 nM | 1000 nM |
|---|---|---|---|---|---|---|
| $<I>$[3)] | 38.0 ± 3.8 photons/ms | 33.2 ± 3.3 photons/ms | 31.3 ± 3.1 photons/ms | 11.1 ± 1.1 photons/ms | 10.3 ± 1.0 photons/ms | 9.09 ± 0.91 photons/ms |
| $b$[4)] | 14.6 ± 1.5 photons/ms | 14.6 photons/ms | 14.6 photons/ms | 14.6 photons/ms | 14.6 photons/ms | 14.6 photons/ms |
| $\tau_R$[5)] | 352 ± 2.2 μs | 356 ± 2.2 μs | 364 ± 2.6 μs | 1180 ± 5.9 μs | 1070 ± 6.2 μs | 1100 ± 6.6 μs |
| $n_R$[6)] | 1.58 ± 0.02 | 1.39 ± 0.19 | 1.42 ± 0.17 | 0.45 ± 0.06 | 1.13 ± 0.10 | 1.24 ± 0.11 |
| $a$[7)] | 1.00 | 0.98 ± 0.16 | 0.94 ± 0.15 | 1.00 ± 0.17 | 0.43 ± 0.06 | 0.35 ± 0.05 |
| $\tau_A$[8)] | 79 μs | 79 μs | 79 μs | 79 μs | 79 μs | 79 μs |
| $n_A$[9)] | 1.02 ± 0.37 | 0.91 ± 0.37 | 0.80 ± 0.34 | 0.31 ± 0.12 | 0.21 ± 0.11 | 0.19 ± 0.10 |
| $s$[10)] | 8.0 | 8.0 | 8.0 | 8.0 | 8.0 | 8.0 |
| $R_H$[11)] | 2.73 ± 0.23 nm | 2.75 ± 0.24 nm | 2.81 ± 0.24 nm | 9.10 ± 0.78 nm | 8.28 ± 0.71 nm | 8.54 ± 0.73 nm |

[1)]The FCS data obtained for 488-rA$_{30}$ excited at 484 nm was analyzed based on eqs. S1 ~ S3. The definitions of the parameters were explained in Supporting Texts. The FCS data and fitted curves were presented in Supporting Figure S2B. The errors of the parameters were estimated by the method explained in Supporting Texts. [2)]The N protein concentration. [3)]The averaged fluorescence intensity during the FCS measurements used for the fitting. The error ranges were roughly estimated as 10 % of the measured fluorescence intensity. [4)]The brightness of the free Alexa. The value estimated for the data in the absence of the N protein assuming $a = 1$ was used as the fixed fitting parameter for the data in the presence of the N protein. [5)]The translational diffusion time for the labeled RNA obtained by the fitting. [6)]The number of the labeled RNA in the observation volume obtained by the fitting. [7)]The ratio of the brightness of the labeled RNA relative to that of the free Alexa. The value was fixed to 1 for the data in the absence of the N protein to estimate $b$, and set as a fitting parameter for the data in the presence of the N protein. [8)]The translational diffusion time for the free Alexa. The value was fixed to the value determined separately. [9)]The number of the free Alexa in the observation volume estimated using eq. S2 and other fitted parameters. [10)]The ratio of the axial radius to the radial radius of the observation volume. The value was fixed to the value determined separately. [11)]The hydrodynamic radius of the labeled RNA estimated from $\tau_R$ using eqs. S4 and S5.

**Supporting Table S1C. The parameters used in and obtained by the fitting analysis of the FCS data for 488-rA$_{20}$ excited at 484 nm**[1)]

| Concentration[2)] | 0 nM | 0.1 nM | 1 nM | 10 nM | 100 nM | 1000 nM |
|---|---|---|---|---|---|---|
| $<I>$[3)] | 38.0 ± 3.8 photons/ms | 41.3 ± 4.1 photons/ms | 34.2 ± 3.4 photons/ms | 24.3 ± 2.4 photons/ms | 9.74 ± 0.97 photons/ms | 9.83 ± 0.98 photons/ms |
| $b$[4)] | 14.9 ± 1.5 photons/ms | 14.9 photons/ms | 14.9 photons/ms | 14.9 photons/ms | 14.9 photons/ms | 14.9 photons/ms |
| $\tau_R$[5)] | 299 ± 1.8 μs | 303 ± 1.8 μs | 366 ± 2.1 μs | 764 ± 6.6 μs | 1140 ± 4.0 μs | 1070 ± 5.4 μs |
| $n_R$[6)] | 1.52 ± 0.02 | 1.53 ± 0.23 | 1.10 ± 0.22 | 0.57 ± 0.18 | 0.95 ± 0.08 | 1.27 ± 0.08 |
| $a$[7)] | 1.00 | 1.04 ± 0.18 | 1.05 ± 0.21 | 1.12 ± 0.29 | 0.49 ± 0.07 | 0.39 ± 0.05 |
| $\tau_A$[8)] | 79 μs | 79 μs | 79 μs | 79 μs | 79 μs | 79 μs |
| $n_A$[9)] | 1.04 ± 0.36 | 1.19 ± 0.46 | 1.14 ± 0.40 | 1.00 ± 0.31 | 0.19 ± 0.10 | 0.16 ± 0.10 |
| $s$[10)] | 7.1 | 7.1 | 7.1 | 7.1 | 7.1 | 7.1 |
| $R_H$[11)] | 2.32 ± 0.20 nm | 2.35 ± 0.20 nm | 2.84 ± 0.24 nm | 5.91 ± 0.51 nm | 8.83 ± 0.75 nm | 8.26 ± 0.70 nm |

[1)]The FCS data obtained for 488-rA$_{20}$ excited at 484 nm was analyzed based on eqs. S1 ~ S3. The definitions of the parameters were explained in Supporting Texts. The FCS data and fitted curves were presented in Supporting Figure S2C. The errors of the parameters were estimated by the method explained in Supporting Texts. [2)]The N protein concentration. [3)]The averaged fluorescence intensity during the FCS measurements used for the fitting. The error ranges were roughly estimated as 10 % of the measured fluorescence intensity. [4)]The brightness of the free Alexa. The value estimated for the data in the absence of the N protein assuming $a = 1$ was used as the fixed fitting parameter for the data in the presence of the N protein. [5)]The translational diffusion time for the labeled RNA obtained by the fitting. [6)]The number of the labeled RNA in the observation volume obtained by the fitting. [7)]The ratio of the brightness of the labeled RNA relative to that of the free Alexa. The value was fixed to 1 for the data in the absence of the N protein to estimate $b$, and set as a fitting parameter for the data in the presence of the N protein. [8)]The translational diffusion time for the free Alexa. The value was fixed to the value determined separately. [9)]The number of the free Alexa in the observation volume estimated using eq. S2 and other fitted parameters. [10)]The ratio of the axial radius to the radial radius of the observation volume. The value was fixed to the value determined separately. [11)]The hydrodynamic radius of the labeled RNA estimated from $\tau_R$ using eqs. S4 and S5.

**Supporting Table S1D. The parameters used in and obtained by the fitting analysis of the FCS data for 488-SL4 excited at 484 nm**[1)]

| Concentration[2)] | 0 nM | 0.1 nM | 1 nM | 10 nM | 100 nM | 1000 nM |
|---|---|---|---|---|---|---|
| $<I>$[3)] | 28.3 ± 2.8 photons/ms | 12.6 ± 1.3 photons/ms | 19.9 ± 2.0 photons/ms | 11.9 ± 1.2 photons/ms | 10.6 ± 1.1 photons/ms | 12.6 ± 1.3 photons/ms |
| $b$[4)] | 20.1 ± 2.0 photons/ms | 20.1 photons/ms | 20.1 photons/ms | 20.1 photons/ms | 20.1 photons/ms | 20.1 photons/ms |
| $\tau_R$[5)] | 391 ± 2.3 μs | 453 ± 2.4 μs | 643 ± 5.9 μs | 1330 ± 8.0 μs | 1130 ± 6.5 μs | 1160 ± 6.2 μs |
| $n_R$[6)] | 0.84 ± 0.01 | 0.59 ± 0.06 | 0.56 ± 0.10 | 0.48 ± 0.05 | 1.15 ± 0.08 | 1.47 ± 0.10 |
| $a$[7)] | 1.00 | 0.69 ± 0.11 | 0.96 ± 0.18 | 0.80 ± 0.12 | 0.34 ± 0.05 | 0.32 ± 0.04 |
| $\tau_A$[8)] | 79 μs | 79 μs | 79 μs | 79 μs | 79 μs | 79 μs |
| $n_A$[9)] | 0.58 ± 0.20 | 0.22 ± 0.10 | 0.45 ± 0.17 | 0.21 ± 0.09 | 0.13 ± 0.08 | 0.16 ± 0.09 |
| $s$[10)] | 8.4 | 8.4 | 8.4 | 8.4 | 8.4 | 8.4 |
| $R_H$[11)] | 3.03 ± 0.26 nm | 3.51 ± 0.30 nm | 4.90 ± 0.43 nm | 10.32 ± 0.88 nm | 8.71 ± 0.74 nm | 8.95 ± 0.76 nm |

[1)]The FCS data obtained for 488-SL4 excited at 484 nm was analyzed based on eqs. S1 ~ S3. The definitions of the parameters were explained in Supporting Texts. The FCS data and fitted curves were presented in Supporting Figure S2D. The errors of the parameters were estimated by the method explained in Supporting Texts. [2)]The N protein concentration. [3)]The averaged fluorescence intensity during the FCS measurements used for the fitting. The error ranges were roughly estimated as 10 % of the measured fluorescence intensity. [4)]The brightness of the free Alexa. The value estimated for the data in the absence of the N protein assuming $a = 1$ was used as the fixed fitting parameter for the data in the presence of the N protein. [5)]The translational diffusion time for the labeled RNA obtained by the fitting. [6)]The number of the labeled RNA in the observation volume obtained by the fitting. [7)]The ratio of the brightness of the labeled RNA relative to that of the free Alexa. The value was fixed to 1 for the data in the absence of the N protein to estimate $b$, and set as a fitting parameter for the data in the presence of the N protein. [8)]The translational diffusion time for the free Alexa. The value was fixed to the value determined separately. [9)]The number of the free Alexa in the observation volume estimated using eq. S2 and other fitted parameters. [10)]The ratio of the axial radius to the radial radius of the observation volume. The value was fixed to the value determined separately. [11)]The hydrodynamic radius of the labeled RNA estimated from $\tau_R$ using eqs. S4 and S5.

**Supporting Table S2A. The parameters used in and obtained by the fitting analysis of the FCS data for 488-r$A_{40}$-647 excited at 642 nm[1)]**

| Concentration[2)] | 0 nM | 10 nM | 100 nM |
|---|---|---|---|
| $<I>$[3)] | 55.1 ± 5.5 photons/ms | 35.0 ± 3.5 photons/ms | 28.1 ± 2.8 photons/ms |
| $b$[4)] | 22.4 ± 2.2 photons/ms | 22.4 photons/ms | 22.4 photons/ms |
| $\tau_R$[5)] | 975 ± 29 μs | 2660 ± 60 μs | 2670 ± 61 μs |
| $n_R$[6)] | 1.20 ± 0.04 | 0.44 ± 0.05 | 0.65 ± 0.06 |
| $a$[7)] | 1.00 | 2.23 ± 0.36 | 1.31 ± 0.19 |
| $\tau_A$[8)] | 198 μs | 198 μs | 198 μs |
| $n_A$[9)] | 1.25 ± 0.35 | 0.58 ± 0.25 | 0.40 ± 0.19 |
| $s$[10)] | 11.2 | 11.2 | 11.2 |
| $R_H$[11)] | 7.55 ± 0.68 nm | 20.61 ± 1.82 nm | 20.67 ± 1.82 nm |

[1)]The FCS data obtained for 488-r$A_{40}$-647 excited at 642 nm was analyzed based on eqs. S1 ~ S3. The definitions of the parameters were explained in Supporting Texts. The FCS data and fitted curves were presented in Supporting Figure S3A. The errors of the parameters were estimated by the method explained in Supporting Texts. [2)]The N protein concentration. [3)]The averaged fluorescence intensity during the FCS measurements used for the fitting. The error ranges were roughly estimated as 10 % of the measured fluorescence intensity. [4)]The brightness of the free Alexa. The value estimated for the data in the absence of the N protein assuming $a = 1$ was used as the fixed fitting parameter for the data in the presence of the N protein. [5)]The translational diffusion time for the labeled RNA obtained by the fitting. [6)]The number of the labeled RNA in the observation volume obtained by the fitting. [7)]The ratio of the brightness of the labeled RNA relative to that of the free Alexa. The value was fixed to 1 for the data in the absence of the N protein to estimate $b$, and set as a fitting parameter for the data in the presence of the N protein. [8)]The translational diffusion time for the free Alexa. The value was fixed to the value determined separately. [9)]The number of the free Alexa in the observation volume estimated using eq. S2 and other fitted parameters. [10)]The ratio of the axial radius to the radial radius of the observation volume. The value was fixed to the value determined separately. [11)]The hydrodynamic radius of the labeled RNA estimated from $\tau_R$ using eqs. S4 and S5.

**Supporting Table S2B. The parameters used in and obtained by the fitting analysis of the FCS data for 488-SL4-647 excited at 642 nm[1)]**

| Concentration[2)] | 0 nM | 10 nM | 100 nM |
|---|---|---|---|
| $<I>$[3)] | 131 ± 13 photons/ms | 57.3 ± 5.7 photons/ms | 90.1 ± 9.0 photons/ms |
| $b$[4)] | 19.9 ± 2.0 photons/ms | 19.9 photons/ms | 19.9 photons/ms |
| $\tau_R$[5)] | 945 ± 22 μs | 2710 ± 61 μs | 2830 ± 62 μs |
| $n_R$[6)] | 2.88 ± 0.08 | 0.25 ± 0.06 | 1.05 ± 0.16 |
| $a$[7)] | 1.00 | 5.65 ± 1.2 | 2.49 ± 0.44 |
| $\tau_A$[8)] | 195 μs | 195 μs | 195 μs |
| $n_A$[9)] | 3.70 ± 0.93 | 1.49 ± 0.52 | 1.93 ± 0.75 |
| $s$[10)] | 11.0 | 11.0 | 11.0 |
| $R_H$[11)] | 7.32 ± 0.64 nm | 21.0 ± 1.9 nm | 21.9 ± 1.9 nm |

[1)]The FCS data obtained for 488-SL4-647 excited at 642 nm was analyzed based on eqs. S1 ~ S3. The definitions of the parameters were explained in Supporting Texts. The FCS data and fitted curves were presented in Supporting Figure S3A. The errors of the parameters were estimated by the method explained in Supporting Texts. [2)]The N protein concentration. [3)]The averaged fluorescence intensity during the FCS measurements used for the fitting. The error ranges were roughly estimated as 10 % of the measured fluorescence intensity. [4)]The brightness of the free Alexa. The value estimated for the data in the absence of the N protein assuming $a = 1$ was used as the fixed fitting parameter for the data in the presence of the N protein. [5)]The translational diffusion time for the labeled RNA obtained by the fitting. [6)]The number of the labeled RNA in the observation volume obtained by the fitting. [7)]The ratio of the brightness of the labeled RNA relative to that of the free Alexa. The value was fixed to 1 for the data in the absence of the N protein to estimate $b$, and set as a fitting parameter for the data in the presence of the N protein. [8)]The translational diffusion time for the free Alexa. The value was fixed to the value determined separately. [9)]The number of the free Alexa in the observation volume estimated using eq. S2 and other fitted parameters. [10)]The ratio of the axial radius to the radial radius of the observation volume. The value was fixed to the value determined separately. [11)]The hydrodynamic radius of the labeled RNA estimated from $\tau_R$ using eqs. S4 and S5.
